# Therapy-induced senescence upregulates antigen presentation machinery and triggers anti-tumor immunity in Acute Myeloid Leukemia

**DOI:** 10.1101/2022.11.17.515658

**Authors:** Diego Gilioli, Simona Fusco, Teresa Tavella, Kety Giannetti, Anastasia Conti, Antonella Santoro, Edoardo Carsana, Stefano Beretta, Martin Schönlein, Valentina Gambacorta, Federico Mario Aletti, Matteo Giovanni Carrabba, Chiara Bonini, Fabio Ciceri, Ivan Merelli, Luca Vago, Clemens Schmitt, Raffaella Di Micco

**Affiliations:** Unit of Senescence in Stem Cell Aging, Differentiation and Cancer, San Raffaele Telethon Institute for Gene Therapy (SR-TIGET), IRCCS San Raffaele Scientific Institute, Milano, Italy; Unit of Immunogenetics, Leukemia Genomics and Immunobiology, Division of Immunology, Transplantation and Infectious Disease, IRCCS San Raffaele Scientific Institute, Milano, Italy; Institute for Biomedical Technologies, National Research Council, Segrate, Italy; San Raffaele Telethon Institute for Gene Therapy (SR-TIGET), IRCCS San Raffaele Scientific Institute, Milano, Italy; Medical Department of Hematology, Oncology and Tumor Immunology, University Medical Center Berlin, Germany; Department of Oncology, Hematology and Bone Marrow Transplantation With Section of Pneumology, University Medical Center Hamburg-Eppendorf, Hamburg, Germany; Hematology and Bone Marrow Transplantation Unit, IRCCS San Raffaele Scientific Institute, Milano, Italy; Unit of Experimental Hematology, Division of Immunology, Transplantation and Infectious Disease, IRCCS San Raffaele Scientific Institute, Milano, Italy; Vita-Salute San Raffaele University, Milano, Italy; Molecular Cancer Research Center, Charité, Berlin

## Abstract

Acute myeloid leukemia (AML) is an aggressive hematological malignancy often curable only by using intensive chemotherapy. Nonetheless, resistance/early relapses are frequent, underscoring the need to investigate the molecular events occurring shortly after chemotherapy. Therapy-induced senescence (TIS) is a fail-safe tumor suppressive mechanism that may elicit immune-mediated responses contributing to senescent cell clearance. Yet, TIS functional role in AML eradication and immune surveillance early post-chemotherapy remains ill-defined. By combining transcriptional and cellular-based evaluation of senescence markers in AML patient samples, we found upregulation of senescence-associated genes and interferon gene categories with concomitant induction of HLA class I and class II molecules, pointing to a causal link between TIS and leukemia immunogenicity. Consistently, senescence-competent AML samples activated autologous CD4^+^ and CD8^+^ T cells and improved leukemia recognition by both T-cell subsets. Lastly, the anti-leukemic activity of Immune Checkpoint Blockades (ICBs) was enhanced upon senescence engagement in AML. Altogether, our results identify senescence as a potent immune-related anti-leukemic mechanism that may rapidly translate into innovative senescence-based strategies to prevent AML relapse.

**STATEMENT OF SIGNIFICANCE:** Our findings uncover a novel link between senescence induction and leukemia immune recognition by T cells via upregulation of antigen presentation machinery components, providing the basis for conceptually novel senescence-based targeted immunotherapeutic regimens for AML patients.

## INTRODUCTION

Acute myeloid leukemia (AML) is an aggressive disease characterized by aberrant proliferation of myeloid progenitors in the bone marrow (BM). Currently, for many AML patients first-line treatment leverages on intensive chemotherapy based on the combination of cytosine arabinoside (ARA-C) and anthracyclines^1^. However, treatment failure frequently occurs^2^, thus highlighting the need to delve deeper into the molecular events occurring early upon chemotherapy to inform better therapeutic strategies. Cancer cells respond to anticancer therapies by either activating an apoptotic program or by entering a state of permanent cell cycle arrest known as Therapy-Induced Senescence (TIS), characterized by protracted DNA Damage Response (DDR), enhanced expression of Senescence Associated-β-galactosidase (SA-β-GAL) and widespread activation of pro- and anti-inflammatory molecules, growth factors and metalloproteases, collectively named as Senescence Associated Secretory Phenotype (SASP)^3–6^. Through the SASP, senescent cells communicate with bystander cells and if persisting, generate a milieu that may even favor the emergence of more aggressive clones contributing to therapy resistance in the long-term^7,8^. Furthermore, senescent cells may spontaneously exit their proliferation block and be reprogrammed into more stem-like cells to promote chemoresistance and relapse^9^. Despite the evidence of long-term detrimental effects of senescence in cancer, TIS cells can also engage in crosstalks with the host immune system, ultimately resulting in their immune-mediated clearance^10^. Specifically, senescent cells were shown to upregulate activating ligands on their surface and favor the recruitment and activation of several cell types of the innate immune system, including macrophages^11,12^, natural killer^13^, dendritic and invariant natural killer T cells^14^, thus contributing to effective senescence immune-surveillance in relevant mouse models of replicative and oncogene-induced senescence. When activated, cells of the innate immune system may in turn initiate and potentiate adaptive immunity against senescent cells, unleashing mainly cytotoxic CD8^+^ T cell responses^15,16^, while the contribution of the CD4^+^ T cells to senescence eradication was, to date, only limited to models of premalignant senescent hepatocytes upon MHC class II upregulation^17^.

Here, we investigated the biological function of senescence induction early upon chemotherapy in patient-derived AML samples and dissected the interplay between TIS and leukemia immune surveillance by the adaptive immune system. Our findings indicate that senescence-inducing therapies may be clinically exploited to potentiate anti-leukemic immune responses as monotherapy or in combination with immune-checkpoint blockades.

## RESULTS

### *Ex-vivo* chemotherapy triggers a senescence-associated transcriptional and cellular program in primary AML blasts

To gain insights into the molecular events triggered by chemotherapy in AML, we analyzed peripheral blood and bone marrow samples from a cohort of 21 AML patients bearing different leukemia-associated genetic alterations (Supplementary Table S1). We first identified 3 patients (UPN03, UPN06 and UPN09) and carried out an exploratory evaluation of the transcriptional consequences triggered early post-chemotherapy by RNA sequencing. Briefly, leukemic cells were positively selected from the bone marrow at diagnosis according to the expression of CD33 or CD34 (Supplementary Figure S1A), then cultured on a layer of irradiated human fibroblasts in presence of a cytokine-rich medium (see Methods Section for details) and treated with cytarabine (ARA-C) for 5 days (Figure 1A). Global transcription profile clearly separated controls from chemotherapy-treated samples as indicated by Principal Component Analysis (PCA) (Supplementary Figure S1B). We first determined the number of Differentially Expressed Genes (DEGs) between the treatment and control groups (adjusted p-value < 0.05) and found 468 DEGs, of which 395 were upregulated and 73 were downregulated (Figure 1B and Supplementary Table S2.1). Interestingly, most DEGs belonged to categories related to DNA damage response, cell cycle and to pro- and anti-inflammatory molecules collectively known as SASP(Figure 1C); of note, *CDKN1A* (p21), a cell cycle inhibitor typically engaged by the senescence program, and several well-characterized SASP genes such as *IL-6, IL1α IL1β*, *CXCL8* and *CCL2* were among the most upregulated upon chemotherapy across all patients analyzed (Figure 1B-C). We next performed Gene Set Enrichment Analysis (GSEA) on gene lists ranked based on log2 fold change, to identify the pathways modulated early after chemotherapy in AML, and found positive and significantly high normalized enrichment score (NES) for genes belonging to categories involved in the DDR (p53 pathway, UV-response-DN), apoptosis, epithelial to mesenchymal transition, metabolism (Pancreas beta cells), inflammation (TNFα signaling via NFkB, IL6 JAK STAT3 signaling, inflammatory response) and interferon responses (Interferon gamma response, Interferon alpha response). Conversely, oxidative phosphorylation, cell cycle regulation (E2F targets, MYC targetsV2), and G2M checkpoint gene categories were negatively enriched (Figure 1D; Supplementary Table S2.2). Furthermore, by comparing our dataset with several published senescent-related gene lists ^8,18–24^, we confirmed a significant transcriptional induction of senescence-related genes in our cohort of *ex-vivo* ARA-C treated primary AML samples (Figure 1E; Supplementary Table S2.3).

**Figure 1.**
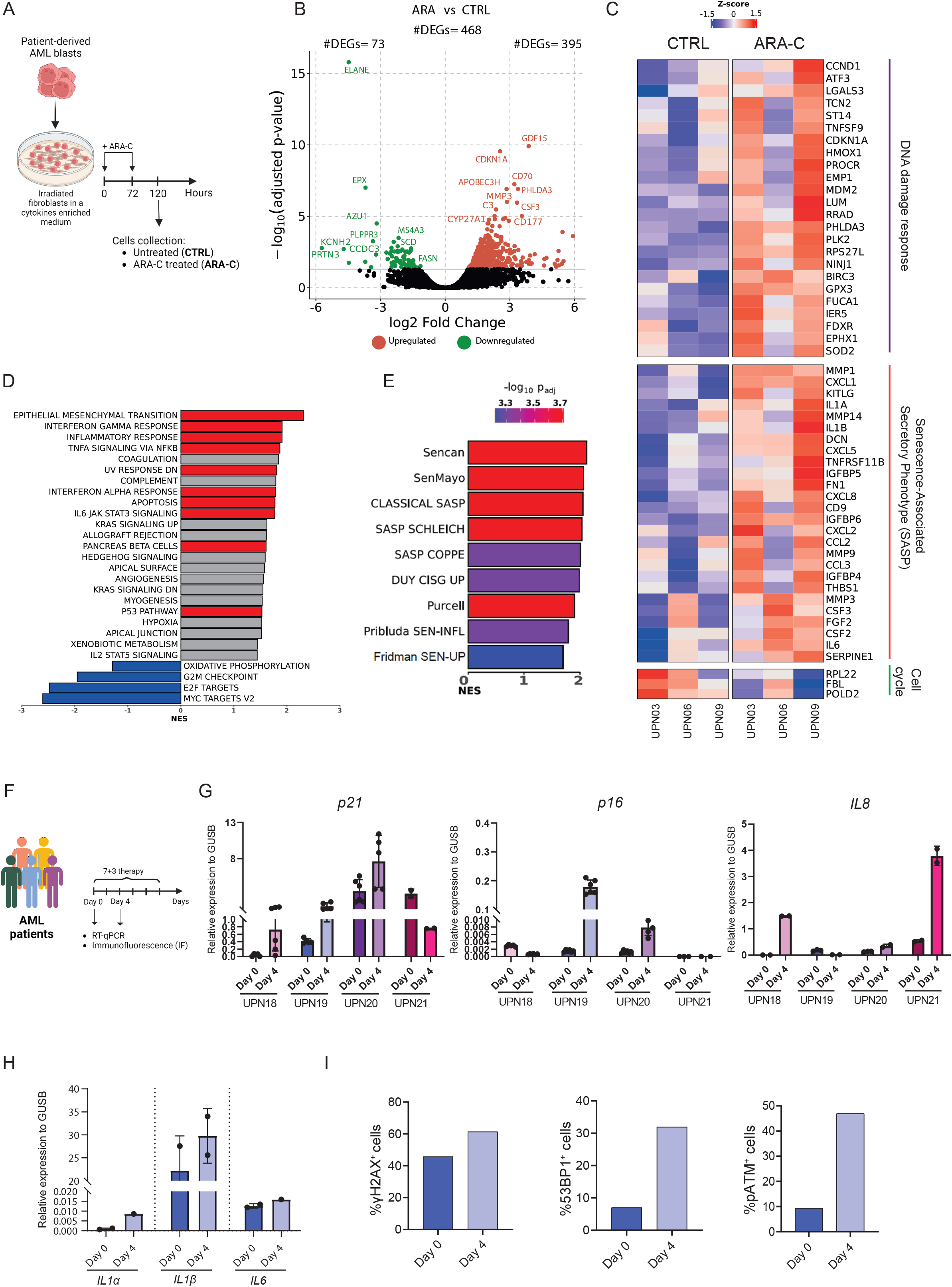
Chemotherapy triggers senescence and inflammatory programs in AML. **A**, Experimental design of *ex-vivo* primary AML treatment. Primary human AML samples were cultured on irradiated human immortalized fibroblasts (BJhTERT) in human bladder carcinoma- (5637) conditioned and cytokine-supplemented medium. After 3 days, blasts purification was performed according to leukemia-associated immunophenotype (CD34 or CD33 positivity within hCD45dim cells, see Supplementary Figure S1A for gating strategy). ARA-C was added twice (at times 0 and at 72 hours) at a 410 nM final concentration. After 5 days of culture (120 hours), cells were collected for RNA-seq analysis. **B**, Volcano plot representing differential expression (log_2_ Fold Change) vs. significance (-log_10_(adjusted p-value)) of gene expression analysis results between CTRL and ARA-C treated AML samples. Of 468 Differentially Expressed Genes (DEGs), 395 were upregulated (red) and 73 were downregulated (green). Top 10 upregulated and downregulated genes are indicated. **C**, Heatmap depicting the normalized (regularized log_2_ abundance, *rlog*) and scaled expression of a subset of DEGs from the contrast ARA-C vs CTRL. Genes involved in DNA damage response, Senescence Associated Secretory Phenotype (SASP) and cell cycle are indicated. Reported SASP genes were collected from the following published signatures (SASP SCHLEICH, CLASSICAL SASP, SASP COPPE). **D**, Bar plot showing the Normalized Enriched Score (NES) of significant categories from the Gene set enrichment analysis (GSEA). Reported categories are from the hallmark v 2.7 gene list. In red and blue the categories described in maintext. **E**, GSEA results showing positive NES of significant senescence signatures. **F**, Experimental design of the clinical protocol for chemotherapy treatment in AML patients. Samples were collected from AML patients at Diagnosis (Day 0) or 4 days after treatment started (Day 4). **G**, Gene expression analysis of *p21, p16* and *IL8* by RT-qPCR on AML blasts purified from 4 AML patients (UPN18, UPN19, UPN20, UPN21) at Day 0 and Day 4. Each dot indicates technical replicates. *GUSB* was used as a housekeeping gene. **H**, Gene expression analysis of *IL1α, IL1β and IL6* by RT-qPCR on AML blasts purified from UPN21 at Day 0 and Day 4. Each dot indicates technical replicates. *GUSB* was used as a housekeeping gene. **I**, Quantification of positive cells for γH2AX, 53BP1 and pATM-S1981 (pATM) on AML blasts purified from UPN21 at Day 0 and Day 4 detected by immunofluorescence analysis. Up to 300 nuclei were analyzed.

As most patients were wildtype for TP53, to assess whether a functional p53 is required for the observed transcriptional programs engaged by chemotherapy, we took advantage of 2 leukemic cell lines, namely THP1 and OCI-AML3, with respectively inactive (26-base deletion beginning at codon 174 of the p53 coding sequence) or active (WT) p53. Upon 7 days of ARA-C treatment (Supplementary Figure S2A) we found a persistent proliferation arrest in both cell lines (Supplementary Figure S2B-C), accompanied by either very low (THP-1, Supplementary Figure S2D) or modest (OCI-AML3, Supplementary Figure S2E) apoptosis induction. This was associated with the up-regulation of p21 at the protein level (Supplementary Figure S2F), up-regulation of SASP genes (Supplementary Figure S2G-H) and increased SA-β-GAL activity (Supplementary Figure S2I-K), despite the presence or absence of functional p53 (Supplementary Figure S2F), suggesting that such mechanism may be conserved even in AML patients with inactive TP53. In line with this, the only sample with TP53 mutation in our cohort was capable to activate senescence upon chemotherapy (Supplementary Table S1).

We next analyzed the effects of induction chemotherapy in freshly harvested *ex vivo* samples, collected from 4 patients (UPN18, UPN19, UPN20 and UPN21) before the start (Day 0) and during (Day 4) induction chemotherapy (Supplementary Table S1) (Figure 1F). Despite the expected inter-patient variability, we consistently observed therapy-related activation of *CDKN1A* (p21), often combined with *CDKN2A* (p16) and/or *IL8* upregulation (Figure 1G). An in-depth characterization of one sample set also revealed upregulation of *IL1α* and *IL1β* (Figure 1H) and widespread accumulation of DDR markers (γH2AX, 53BP1 and pATM) in AML blasts upon *in vivo* chemotherapy (Figure 1I and Supplementary Figure S2L-M).

We then took advantage of both ImageStreamX^25^ and flow cytometry to simultaneously assess and visualize proliferation rates (by EdU incorporation), apoptosis (AnnexinV), and senescence induction (by SA-β-GAL activity) in 17 AML patients at diagnosis and upon *ex-vivo* treatment. As expected, in all samples analyzed chemotherapy treatment resulted in a significant reduction of cell proliferation (Figure 2A), without any overt induction of cell death (Figure 2B). Interestingly, in 10 out of 17 AML primary samples, this was also accompanied by increased levels in SA-β-GAL activity (hereafter defined as “Senescence High” samples). In contrast, 7 out of 17 samples analyzed did not activate senescence based on SA-β-GAL activity upon *ex-vivo* treatment and were thereby defined as “Senescence Low” samples (Figure 2C-E and Supplementary Table S1). Similar to samples exposed to chemotherapy *in vitro*, we found that SA-β-GAL activity increased in AML collected from patients on day 4 of chemotherapy (Figure 2F-G).

**Figure 2.**
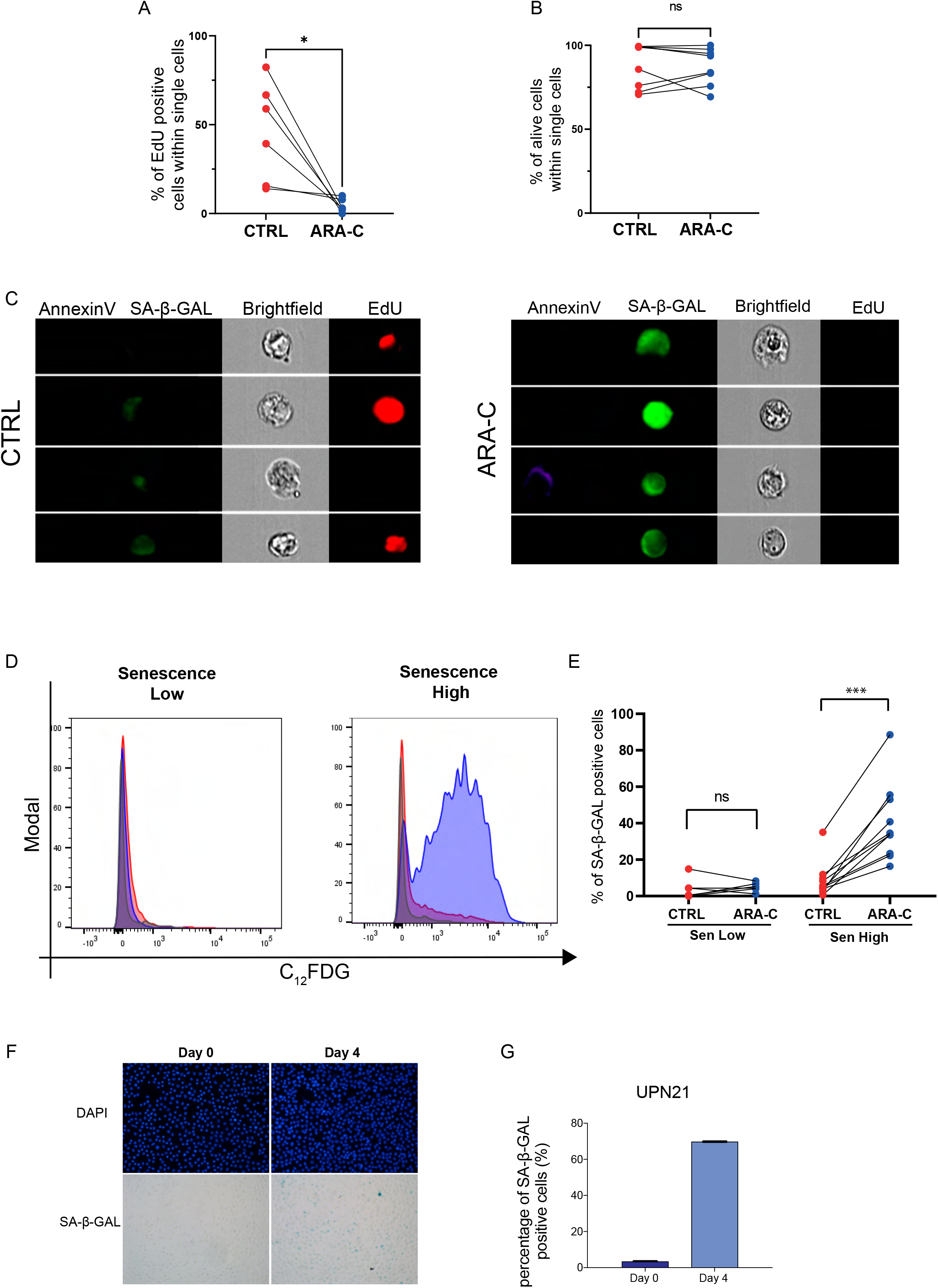
SA-β-Gal levels stratify AML patients in “Senescence High” and “Senescence Low” groups. **A**, Proliferation assay of untreated (CTRL, red) or treated (ARA-C, blue) primary AML samples measured by ImageStreamX. EdU incorporation was used to distinguish cycling (Edu-positive) from non-cycling (Edu-negative) cells. The percentage of cycling cells was plotted among conditions (n=6). Statistical test: Wilcoxon matched-pairs signed rank test. ‘*’ p<0.05 **B**, Viability assay of untreated (CTRL, red) or treated (ARA-C, blue) primary blasts measured by ImageStreamX. The percentage of AnnexinV negative (alive) cells was plotted (n=8). Statistical test: Wilcoxon matched-pairs signed rank test. **C**, Representative images of AnnexinV, Senescence (fluorescent SA-β-GAL), Proliferation (EdU) or brightfield images by ImageStreamX analysis from a representative untreated (CTRL) and ARA-C treated (ARA-C) AML sample. **D**, Distribution of cells along the fluorescent intensity axis, corresponding to SA-β-GAL activity upon C_12_FDG staining (see Methods section). Two representative flow cytometry plots from either “Senescence Low” or “Senescence High” AML patients are shown. **E**, Differential fluorescent SA-β-GAL activity between Senescence Low (left) or Senescence High (right) matched untreated (CTRL, red) and ARA-C (ARA-C, blue) treated primary AML samples (n=17). Statistical test: Wilcoxon matched-pairs signed rank test. ‘***’ p<0.001 **F**, Representative images of DAPI staining (top row) and classic SA-β-GAL staining (bottom row) on AML blasts purified from UPN21 at Day 0 (on the left) and Day 4 (on the right) post-chemotherapy *in vivo*. **G**, Quantification of SA-β-GAL positive cells from panel F.

Altogether, our data indicates that chemotherapy triggers therapy-induced senescence (TIS) in primary AML blasts, albeit to different extent according to SA-β-GAL activity.

### TIS AML upregulates antigen processing and presentation machinery

To dissect the biological processes associated with TIS induction in AML, we functionally annotated upregulated genes obtained by RNA-seq analysis (Figure 1B), with a specific focus on pathways from KEGG, Reactome, and Gene Ontology (GO) databases. Interestingly, we found enriched gene categories related to inflammation, extracellular matrix remodeling and immunity in treated AML samples compared to controls (Figure 3A). When analyzing biological processes (BP), molecular functions (MF), and cellular components (CC) gene set we identified a positive regulation of the adaptive immune system, with several categories involved in leukocyte migration, T cell activation and cytotoxicity, antigen processing and presentation via MHC molecules (Figure 3B, Supplementary Figure S3A). Consistent with this, we found increased expression of Human Leukocyte Antigen (HLA) genes (both class I and II) and their regulators NLR5C (for HLA class I), CD74, and CIITA (for HLA class II) in ARA-C treated blasts compared to controls (Figure 3C). We then evaluated by immunophenotypic analysis the expression levels of both canonical HLA class I (HLA-A, -B, -C,) and HLA class II (HLA-DR) molecules in AML cells treated with ARA-C (Figure 3D-G and Supplementary Figure S3B). Interestingly, we detected upregulation of HLA class I and II molecules mainly in AML samples with more evident induction of senescence genes, henceforth defined “Senescence High” group (Figure 3D-E), and consistently a positive and significant correlation between senescence induction measured by SA-β-GAL and HLA upregulation in ARA-C treated AML blasts (Figure 3F-G).

**Figure 3.**
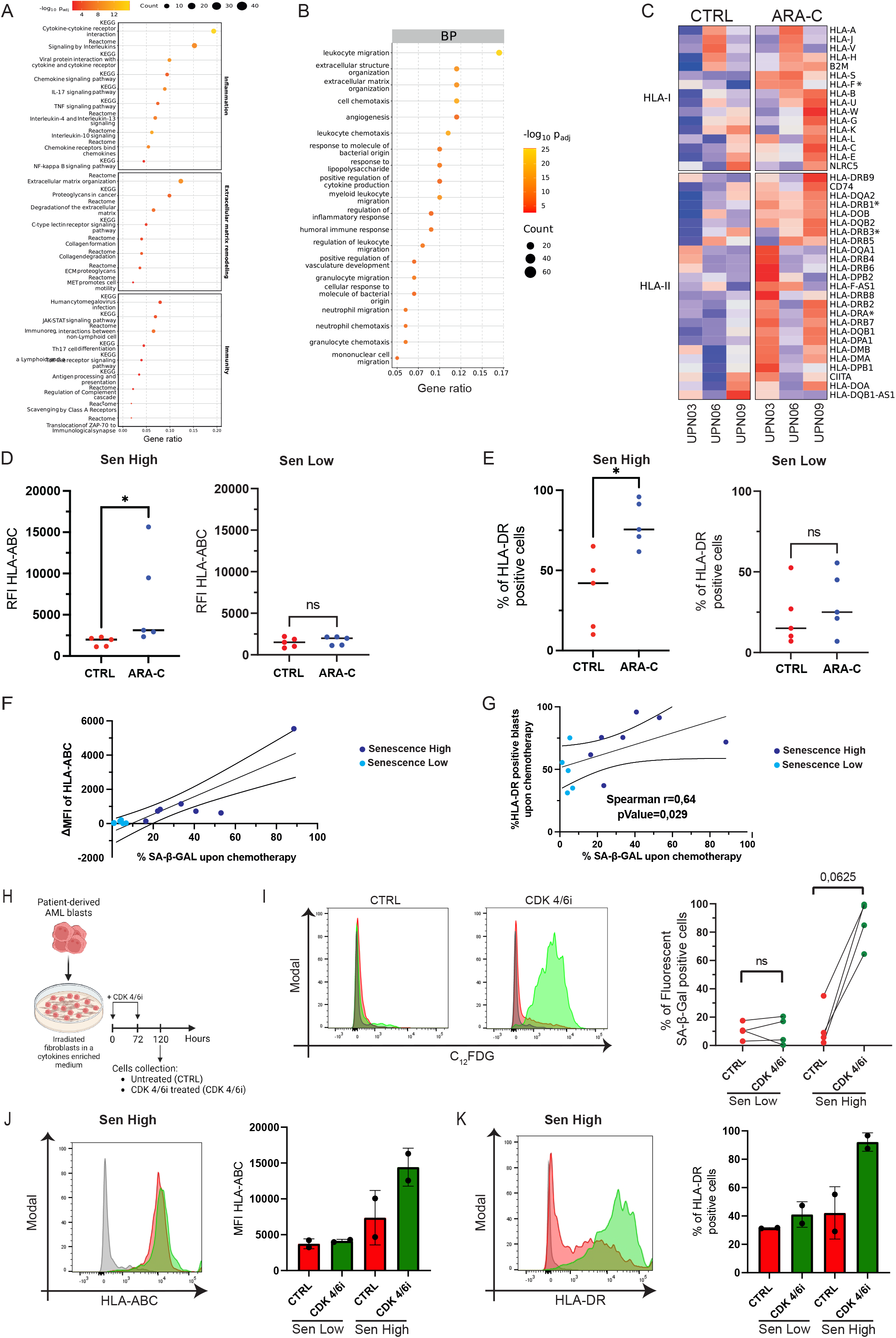
Senescent AML cells upregulate antigen processing and presentation machinery. **A**, Dot plot reporting the results of the Over-Representation Analysis (ORA) computed on positive DEGs. The analysis was conducted on KEGG and Reactome databases. Representative pathways are grouped in manually curated macro-categories: Inflammation, Extracellular matrix remodeling and Immunity. **B**, ORA of positive DEGs conducted on GO terms, reporting selected enriched terms for the biological process (BP) ontology branch. **C**, Heatmap reporting the normalized and scaled expression of HLA genes (class I and II) and their regulators (*NLR5C, CIITA*), for the contrast ARA-C vs CTRL. Asterisks indicate DEGs (FDR < 0.05). **D**, Relative Fluorescence Intensity (RFI) of HLA-ABC as detected by FACS analysis in untreated (CTRL, red) and treated (ARA-C, blue) primary Senescence High (on the left) and Senescence Low (on the right) AML blasts (n=5). Statistical test: Wilcoxon matched-pairs signed rank test. ‘*’ p < 0.05. **E**, Percentage of HLA-DR positive cells as detected by FACS analysis in untreated (CTRL, red) and treated (ARA-C, blue) Senescence High (on the left) and Senescence Low (on the right) primary AML blasts (n=5). Statistical test: Wilcoxon matched-pairs signed rank test. ‘*’p < 0.05. **F,** Correlation between percentage of SA-β-GAL positive cells in ARA-C treated samples (X-axis) and variation of Median Fluorescence Intensity (MFI) of HLA-ABC (class I) signal obtained by CTRL vs. ARA-C treatment comparison (ΔMFI of HLA-ABC, Y-axis); each dot represents a primary AML sample either from the Senescence High (dark blue) or the Senescence Low (light blue) group (n=12). Spearman’s coefficient (r) and p-value are indicated. **G**, Correlation between percentage of SA-β-GAL positive cells of ARA-C treated samples (X-axis) and HLA-DR (class II) positive cells of ARA-C treated samples (Y-axis); each dot represents a primary AML sample either from the Senescence High (dark blue) or the Senescence Low (light blue) group (n=12). Spearman’s coefficient (r) and p-value are indicated**. H**, Experimental design of *ex-vivo* primary AML blasts treatment with CDK 4/6i. CDK 4/6i was added twice (at times 0 and 72 hours) at a 10μM final concentration. After 5 days of culture (120 hours), cells were collected for FACS analyses. **I**, On the left, distribution of cells along the fluorescent intensity axis corresponding to SA-β-Gal activity upon C_12_FDG staining (see Methods section). Here, 2 representative FACS plots with results from either CTRL or CDK 4/6i treated samples are shown. On the right, the percentage of fluorescent SA-β-GAL positive cells from matched untreated (CTRL, red) and CDK 4/6i (CDK 4/6i, green) exposed primary AML blast samples are shown (n=4). Statistical test: Wilcoxon matched-pairs signed rank test. **J**, On the left, a representative image of the distribution of cells along the fluorescent intensity axis corresponding to the HLA-ABC signal; on the right, MFI of HLA-ABC positive cells in untreated (CTRL, red) and treated (CDK 4/6i, green) primary AML blasts from either Senescence Low (left; n=2) and Senescence High (right; n=2) groups. **K**, On the left, a representative image of the distribution of cells along the fluorescent intensity axis corresponding to the HLA-DR signal; on the right, the percentage of HLA-DR positive cells in untreated (CTRL, red) and treated (CDK 4/6i, green) primary AML blasts from either Senescence Low (left; n=2) and Senescence High (right; n=2) group.

We then asked whether senescence establishment *per se*, rather than chemotherapy-induced DNA damage response, would trigger the upregulation of antigen processing and presentation machinery For this purpose, we took advantage of the CDK4/6 inhibitor (CDK4/6i) Palbociclib (Supplementary Figure S4A), a molecule that halts progression from G1 to the S phase of the cell cycle by inhibiting the activity of cyclin D-dependent kinases 4 and 6, and by this inducing senescence^5^. As expected, CDK4/6i treatment arrested the proliferation of AML cell lines preferentially in the G1 phase, rather than in the S phase as for ARA-C treated cells (Supplementary Figure S4B), eliciting a proliferation slowdown in the absence of apoptosis (Supplementary Figure S4C-F), without accumulation of the DDR sensor γH2AX (Supplementary Figure S4G). Accordingly, when measuring SA-β-GAL activity, we confirmed that this treatment triggers senescence in AML blasts (Supplementary Figure S4H). Interestingly, and in line with our model, quantifying HLA class I (Supplementary Figure S4I) and HLA class II (Supplementary Figure S4J) on CDK 4/6i treated cells also revealed an up-regulation similar to that observed in ARA-C treated samples. We next treated primary AML *ex-vivo* blasts with CDK 4/6i for 5 days (Figure 3H) and found an accumulation of SA-β-GAL (Figure 3I) with the concomitant induction of HLA class I and II molecules (Figure 3J-K) specifically in the “Senescence High” group of patients. Importantly, acute treatment with both ARA-C and CDK4/6i was not sufficient to induce senescence (Supplementary Figure S4K), did not result in upregulation of HLA class II (Supplementary Figure S4L), and led to only a modest increase in HLA class I levels for both treated groups compared to control (Supplementary Figure S4M).

Taken together, these data suggest that, independently from the initial trigger (e.g., DNA damage (ARA-C) or cell cycle inhibition (CDK4/6i)), prolonged treatment of AML blasts is required to trigger senescence and up-regulation of HLA molecules and regulators. These results prompted us to investigate the link between senescence induction and activation of the adaptive immune system against senescent cancer cells.

### Therapy-induced senescence promotes T-cell activation

To experimentally assess the immune-related consequences of HLA up-regulation in TIS AML samples, we first co-cultured senescent or control THP-1 cells with CD3^+^ T cells from healthy volunteers and evaluated the activation of both CD4^+^ (Helper) or CD8^+^ (Cytotoxic) T cells. Leukemia-primed T cells were tested against their original targets either left untreated (CTRL) or induced into senescence by ARA-C or CDK4/6i treatment (Figure 4A). T cells co-cultured with IFN-γ treated leukemia cells were used as a positive control^26^. As experimental readouts, we measured T cell proliferation (by label dilution method), activation (by CD25 and CD69 expression levels), and degranulation (by CD107a expression). In all T cell donors tested (n=6), we detected, consistently throughout all read-outs, modest recognition of untreated leukemic blasts and potent increase in T cell activation against leukemia upon induction of senescence by either ARA-C or CDK4/6i treatment. Specifically, we observed a consistent HLA-DR-dependent increase of CD4^+^ T cell proliferation by dilution of CT violet (Supplementary Figure S5A-B; Figure 4B,), paralleled by an increase in activation markers CD69 (Figure 4C, Supplementary Figure S5C) and CD25 (Figure 4D, Supplementary Figure S5D) on the cell surface. When measuring CD8^+^ T cells activation, we found a modest increase in proliferation rates when T cells were tested against senescent or IFN-y treated AML cells compared to controls (Figure 4E); however, a marked increase in degranulation by CD107a expression (Figure 4F, Supplementary Figure S5E) and activation markers CD69 (Figure 4G) and CD25 (Figure 4H) was observed in CD8^+^ T cells when challenged against senescent leukemic cells.

**Figure 4.**
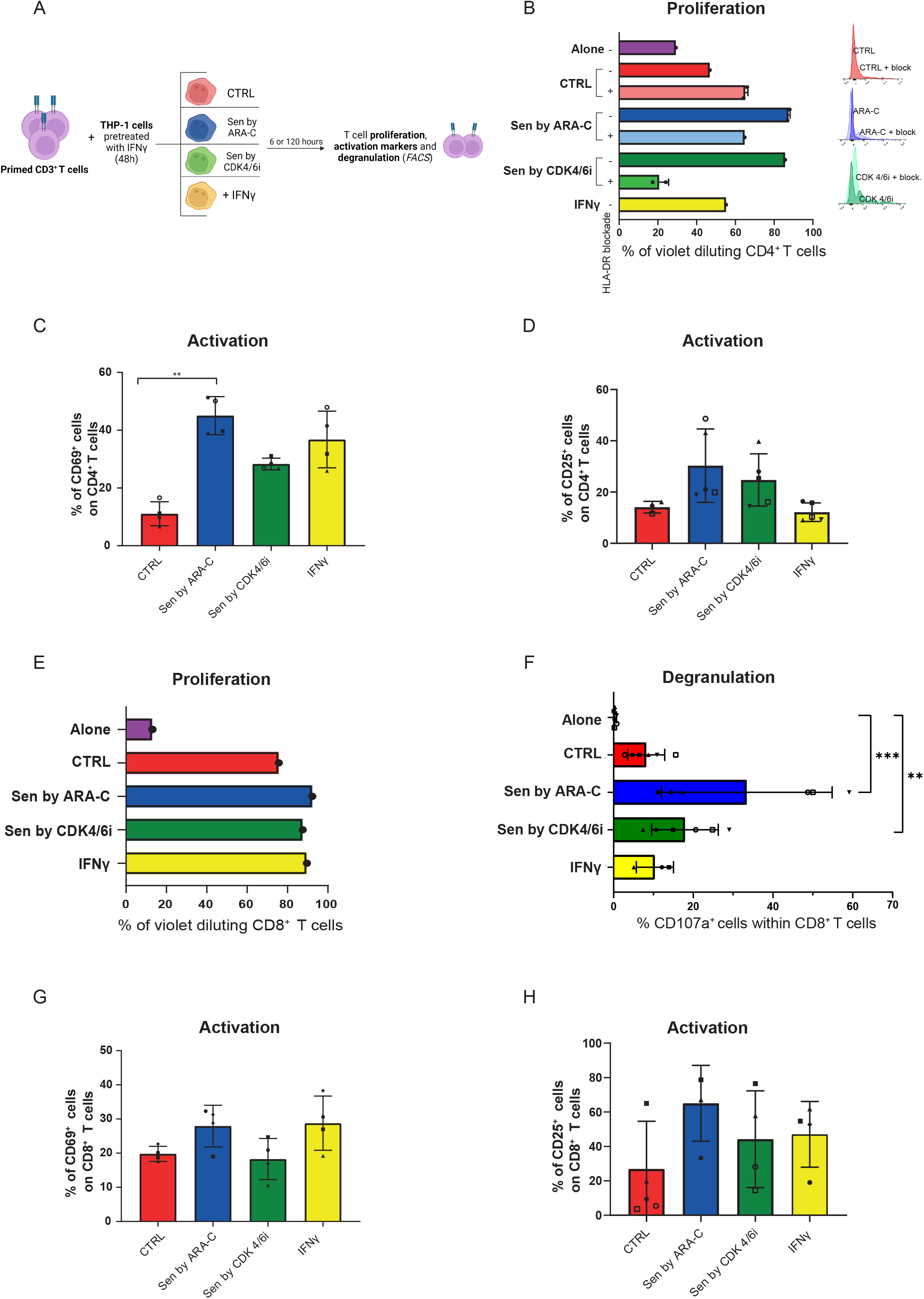
Therapy-induced senescence promotes T-cell activation. **A**, Experimental design of the Mixed Lymphocyte Culture assay. After an initial priming of CD3^+^ T cells by co-culture with 48h-IFNγ-pretreated THP-1 cells, T cells were co-cultured in a 1:1 ratio with THP-1 cells either untreated (CTRL, red) or treated with either ARA-C (Sen by ARA-C, blue; a 7 days treatment was applied) or CDK 4/6i (Sen by CDK 4/6i, green; a 7 days treatment was applied) or IFN-γ (IFN-γ, yellow; a 48h treatment was applied). T cell proliferation and activation were assessed 120 hours post-co-culture. **B**, Proliferating CD4^+^ T cells (measured as % of violet diluting CD4^+^ T cells) 120 hours post *in vitro* culture of T-cells alone (violet bar) or co-cultured with different targets: untreated (CTRL, dark red), treated with ARA-C (Sen by ARA-C, dark blue) or treated with CDK 4/6i (Sen by CDK 4/6i, dark green) or treated with IFN-γ (IFN-γ, yellow). For CTRL (light red), Sen by ARA-C (light blue) or Sen by CDK 4/6i (light green) co-cultures, HLA-DR blockade (through *In Vivo* antihuman HLA-DR MAb 10 μg/ml) was performed; on the right, a representative image of the distribution of cells along the fluorescent intensity axis corresponding to Cell Trace Violet signal, same colors as bars on the left. **C**, Percentage of CD69^+^ cells within the CD4^+^ T cells population, detected by FACS analysis 6 hours after co-culture with different target cells, according to panel A. Each symbol represents different biological replicates (n= 4). Statistical test: Friedman test. ‘**’ p<0.01. **D**, Percentage of CD25^+^ cells within the CD4^+^ T cells population, detected by FACS analysis 120 hours post co-culture after co-culture with different target cells, according to panel A. Each dot represents different biological replicates (n= 3-5). **E**, Proliferating CD8^+^ T cells (measured as % of violet diluting CD8^+^ T cells) 120 hours post *in vitro* culture of T-cells alone (violet) or cocultured with different targets: untreated (CTRL, red), treated with ARA-C (Sen by ARA-C, blue) or treated with CDK 4/6i (Sen by CDK 4/6i, green) or treated with IFN-γ (IFN-γ, yellow). **F**, Quantification by FACS of degranulating CD8^+^ cells (measured as CD107a^+^ cells) 6 hours post CD8^+^ T cells *in vitro* culture alone (violet) or co-cultured with targets as indicated in panel E. Each symbol represents a biological replicate (n= 3-6). Statistical test: Kruskal-Wallis with Dunn’s multiple comparisons. ‘**’p < 0.005, ‘***’p < 0.0005. **G**, Percentage of CD69^+^ cells within the CD8^+^ T cells population, detected by FACS analysis 6 hours after co-culture with different target cells, according to panel A. Each symbol represents a different biological replicate (n=4). **H**, Percentage of CD25^+^ cells within the CD8^+^ T cells population, detected by FACS analysis 120 hours after coculture with different target cells, according to panel A. Each symbol represents distinct T cell biological replicates (n= 3-5).

### Senescence-competent primary AML are sensitized to T cell-mediated immune eradication

We next evaluated the immunogenicity of primary AML samples by taking into account their competence to undergo senescence upon treatment (Senescence High or Senescence Low groups, Figure 2E). We first focused on CD4^+^ T cells, the subset showing higher activation when challenged with senescent AML cells and more consistently implicated in immune-based recognition of leukemia after bone marrow transplantation^26^. As experimental readouts, we assessed T cell proliferation, CD4^+^ T cells and target cells immunological synapse quantification, and their killing activity (Figure 5A). CD4^+^ T cells purified from healthy individuals and primed against primary AML pre-treated with IFN-γ were tested against untreated AML or their ARA-C-treated counterparts. While low and similar levels (approximately 20%) of T cell proliferation rates were observed when T cells were co-cultured with control samples from either Senenescence High or Senescence Low groups, ARA-C treatment boosted CD4^+^ T cell proliferation only when applied to blasts belonging to the Senescence High group (Figure 5B-C), indicating that senescence establishment reinforces T cell activation. To characterize more in depth the physical interactions between senescent AML cells and T cells, we quantified immunological synapse formation, which by stabilizing the adhesion between T and target cells could contribute to T cell-mediated clearance of AML. Exploiting the ImageStreamX technology, we revealed that T cells formed three times more immunological synapses when challenged with senescent blasts (Senescence High) while ARA-C treatment per se without senescence activation in a Senescence Low patient did not lead to these changes (Figure 5D-E, Supplementary Figure S5F). We then evaluated the killing ability of activated T cells against primary AML blasts in real-time by exploiting the IncuCyte live-cell analysis system, combining SA-β-GAL staining (green fluorescence) with a caspase-3 substrate, that emits red fluorescence when a cell undergoes apoptosis (Figure 5F). At the beginning of the co-culture, we observed that AML cells from a Senescence High patient treated with ARA-C displayed a broader area occupied by senescent live cells, positive for the fluorescent SA-β-GAL (green signal) and negative for caspase 3 (red signal) (Figure 5G). Over time, the area occupied by senescent cells decreased to a level comparable to that observed in untreated controls (Figure 5G). Conversely, when analyzing the area occupied by senescent dead cells, which are positive for both green and red signals, we observed that in ARA-C treated cells the area occupied by senescent dead cells increases over time upon co-culture with T cells, suggesting T cell-mediated eradication of senescent blasts (Figure 5H). By contrast, for a primary AML sample belonging to the Senescence Low group, the senescent live cells area occupied by the ARA-C treated cells was very low and comparable to the untreated condition, consistent with its inability to undergo senescence, and both senescence and apoptosis levels were maintained constant over time (Figure 5G, H).

**Figure 5.**
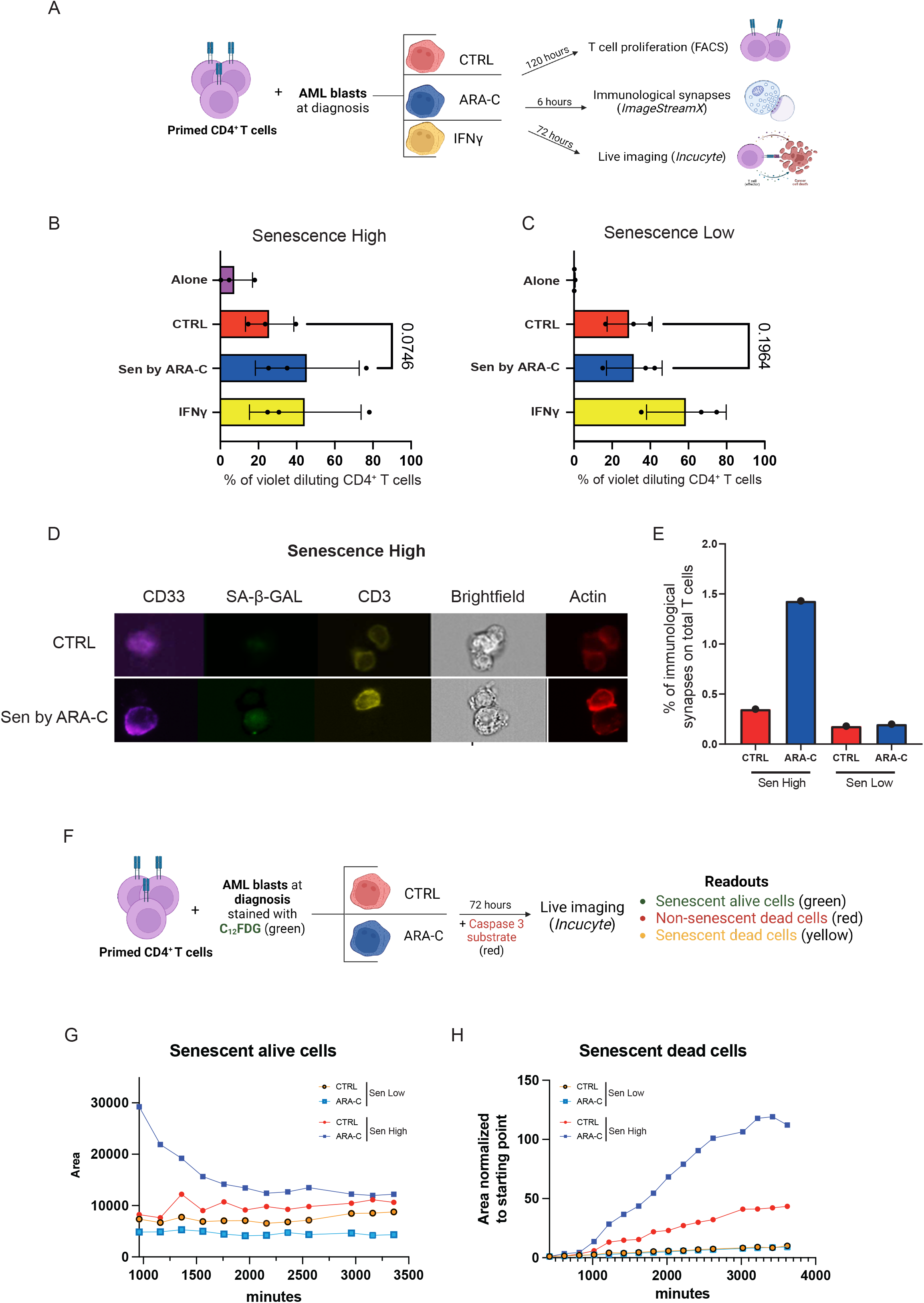
Senescence-competent primary AMLs are sensitized to T cell-mediated immune eradication. **A**, Experimental design of the Mixed Lymphocyte Culture assay. After an initial priming of CD4^+^ T cells by co-culture with 48h-IFNγ-pretreated primary AML samples, T cells were co-cultured in a 1:1 ratio with AML blasts either untreated (CTRL, red) or pre-treated with either ARA-C (ARA-C, blue; a 7 day treatment was applied) or IFN-γ (IFN-γ, yellow; a 48h treatment was applied). Different T cell outcomes were evaluated: proliferation (120 hours post-co-culture), immunological synapses formation (6 hours post co-culture) and killing activity (72 hours post coculture). **B-C,** Proliferating CD4^+^ T cells (measured as % of violet diluting CD4^+^ T cells) 120 hours post *in vitro* culture of T-cells alone (violet) or co-cultured with different targets: untreated (CTRL, red), treated with ARA-C (Sen by ARA-C, blue) or treated with IFN-γ (IFN-γ, yellow) from Senescence high (B) or Senescence low (C) AML patient samples. Each dot represents a biological replicate. Statistical test: paired t-test. **D**, Representative images of the events acquired for evaluation of immunological synapses formation via ImageStreamX. Blasts marker: CD33 (magenta), Senescence: SA-β-GAL (green), T-cell marker: CD3 (yellow), Brightfield and Actin (red). **E**, Percentage of immunological synapses on total T cells co-cultured with either untreated (CTRL, red) or ARA-C treated (ARA-C, blue) primary blasts from a Senescence High (on the left) or a Senescence Low (on the right) patient. **F**, Experimental design of the leukemia killing assay. After priming of CD4^+^ T cells, a 1:1 ratio co-culture with AML blasts incubated with C_12_FDG and either untreated (CTRL, red) or pre-treated with ARA-C (ARA-C, blue; a 7 day treatment was applied). After 72 hours, Caspase 3 substrate is added, then live imaging is performed with Incucyte to detect red and green fluorescence signals giving the following outcomes: Senescent alive cells (green), non-senescent dead cells (red) and senescent dead cells (yellow). **G-H,** Quantification of the area occupied by senescent alive (G) or senescent dead (H) cells over time; blasts from either a Senescence Low (UPN05) or a Senescence High (UPN07) patient either untreated (CTRL) or chemotherapy treated (ARA-C) according to symbol and legend in the figure.

Altogether these findings indicate that senescence establishment after chemotherapy enhances immune-mediated recognition and killing of AML cells by T cells.

### Patient-derived T cells recognize TIS blasts and synergize with immune checkpoint blockades

To confirm our findings on senescence immunogenicity with patient-derived T cells, we performed functional assays on autologous CD3^+^ T cells from the bone marrow of AML patients both at diagnosis (Day 0) or at 30 days from chemotherapy start (Day 30), aiming to assess possible differences in T cell activation states evoked by *in vivo* chemotherapy. We first primed autologous T cells with IFN-γ treated blasts from a Senescent High patient and observed activation of both CD4^+^ and CD8^+^ T cells independently from the treatment applied on blasts, as well as an overall higher activation when T cells harvested at day 30 were used for co-cultures (Fig. 6A-B). We next repeated the co-culture assay by priming T cells from 3 patients belonging to the Senescence High group with autologous ARA-C pre-treated leukemia cells (Figure 6C). In this setting, we observed an increased T cell activation for both subsets (CD4^+^ and CD8^+^) when these were challenged against TIS blasts, and again an overall higher activation level when using T cells harvested at day 30 (Figure 6D). We then challenged autologous T cells from a Senescence Low patient with control or ARA-C treated AML, again primed with ARA-C treated blasts. Interestingly, we did not observe any increase in T cell proliferation after co-culture with treated blasts compared to controls, neither in both T cell subsets nor among T cells collected at the two time points (Figure 6E). When characterizing the T cell compartment by measuring the expression of activation markers, CD69 and CD25, no major changes were found in their levels of expression between the two time points, independently from the capability of AML samples to undergo senescence (Supplementary Figure S6A-B). We also evaluated the combinatorial expression of exhaustion markers TIM-3, LAG-3, and PD-1 by flow cytometry and clustered cells from non-exhausted (green, 0 inhibitory markers) to highly exhausted (red, 3 inhibitory markers) according to the number of markers simultaneously expressed by each individual cell by SPICE^27^. Given that the T cells from the two time points (at diagnosis or day 30) retain a similar activation and exhaustion profile (Supplementary Figure S6C), our data indicate that the enhanced activation observed in T cells at day 30 compared to day 0 against TIS blasts could be ascribed to an expansion of T cells reactive against target blasts, which occurred *in vivo* after chemotherapy treatment.

**Figure 6.**
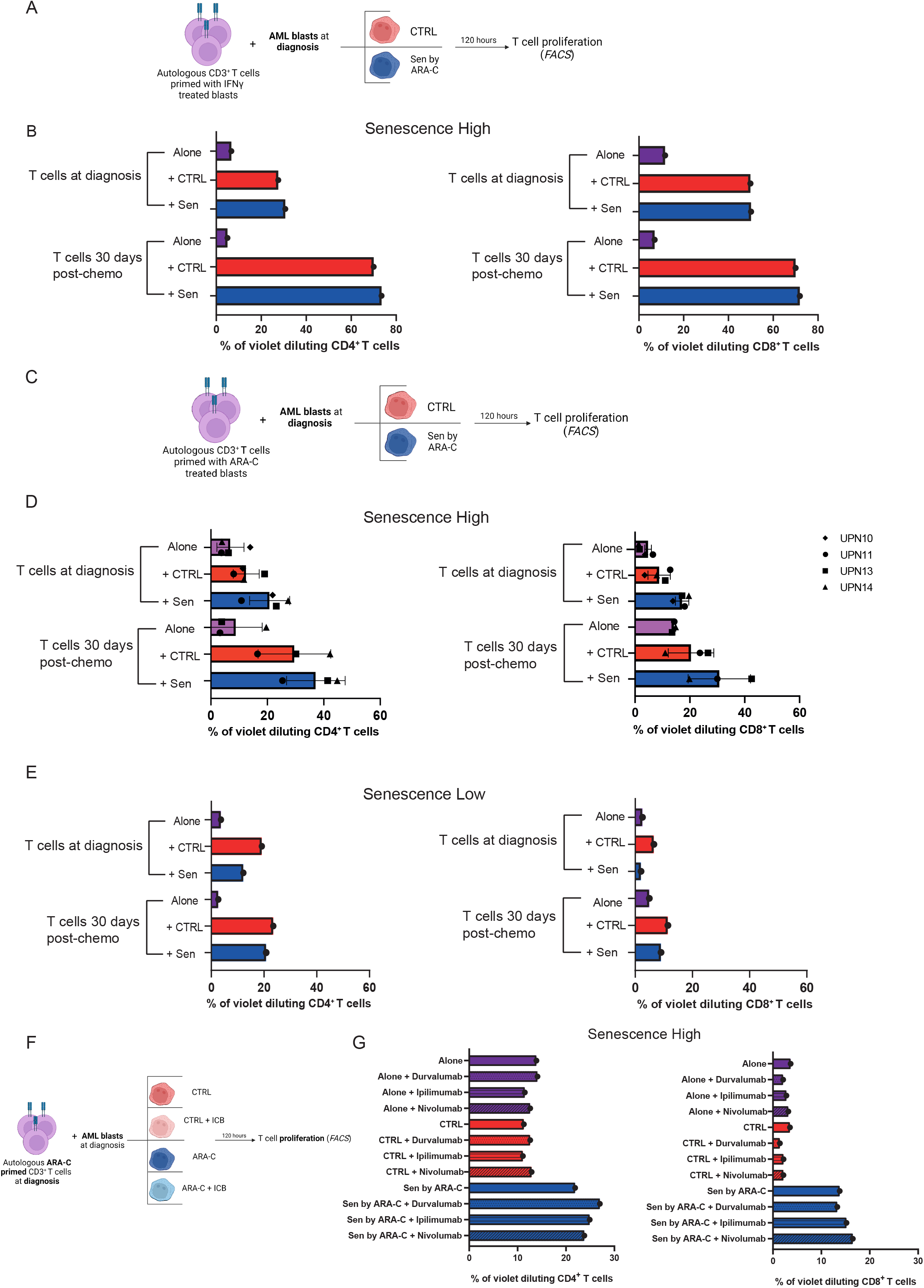
Patient-derived T cells recognize TIS blasts and synergize with immune checkpoint blockades. **A**, Experimental design of the Mixed Lymphocyte Culture assay. After an initial priming of CD3^+^ T cells by co-culture with 48h-IFNγ-pretreated AML blasts, autologous T cells were co-cultured in a 1:1 ratio with AML blasts either untreated (CTRL, red) or pre-treated with ARA-C (ARA-C, blue; a 7 day treatment was applied). T cell proliferation is then assessed 120 hours post-co-culture via FACS analysis. **B**, Proliferating CD4^+^ (on the left) or CD8^+^ (on the right) autologous T cells measured as % of violet diluting T cells, evaluated 120 hours post *in vitro* culture of T-cells alone (violet) or co-cultured with AML blasts from a Senescence High patient (UPN 10) either untreated (CTRL, red) or treated with ARA-C (+Sen, blue). T cells were collected from AML patients either at diagnosis (Day 0) or 30 days post-chemotherapy treatment start (Day 30), as indicated in the graph. **C**, Experimental design of the Mixed Lymphocyte Culture assay. After an initial priming of CD3^+^ T cells by co-culture with ARA-C treated AML blasts, the assay was carried out as indicated in panel A. **D**, Proliferating CD4^+^ (on the left) or CD8^+^ (on the right) autologous T cells measured as % of violet diluting T cells, evaluated 120 hours post *in vitro* culture; see panel B for figure legend. Each dot represents a different biological replicate (patient identifiers are shown as symbols in the figure). All patients analyzed are from the Senescence High group. **E**, Proliferating CD4^+^ (on the left) or CD8^+^ (on the right) autologous T cells measured as % of violet diluting T cells, evaluated 120 hours post *in vitro* culture; see panel B for figure legend. AML blasts from a Senescence Low patient (UPN12) were used for this assay. **F**, Experimental design of the Mixed Lymphocyte Culture assay. After an initial priming of CD3^+^ T cells by coculture with 48h-IFNγ-pretreated AML blasts, autologous T cells were co-cultured in a 1:1 ratio with AML blasts either untreated (CTRL, red) or pre-treated with ARA-C (ARA-C, blue; a 7 day treatment was applied). At the time of co-culture, different ICBs (Ipilimumab, Nivolumab or Durvalumab) were added (light red for CTRL and light blue for ARA-C). T cell proliferation was assessed 120 hours post-co-culture via FACS analysis. **G**, Proliferating CD4^+^ (on the left) or CD8^+^ (on the right) autologous T cells measured as % of violet-diluting T cells, evaluated 120 hours post *in vitro* culture of T-cells alone (violet) or co-cultured with AML blasts from a Senescence High patient (UPN 10) either untreated (CTRL, red) or treated with ARA-C (Sen by ARA-C, blue). T cells were obtained from AML patients at diagnosis.

Immune Checkpoint Blockade (ICB) has shown limited success as monotherapy in AML^28^. Intrigued by the findings that senescence may render AML blasts more immunogenic, we asked whether senescence activation could synergize with ICB treatment, boosting their anti-leukemic potential. We thus tested 3 different ICBs: Ipilimumab (anti-CTLA4), Nivolumab (anti-PD-1), and Durvalumab (anti-PD-L1), adding them to the co-culture between primed autologous CD3^+^T cells and their targets (Figure 6F). When T cells were challenged against ARA-C treated blasts from a Senescence High patient, a modest but consistent increase in T cell proliferation was observed with all ICBs tested in CD4^+^ T cells and only with Nivolumab in CD8^+^ T cells (Figure 6G).

Overall, these data suggest that TIS could be exploited to enhance T cell immune-mediated recognition of leukemic cells in the context of ICBs for AML patients.

## DISCUSSION

Therapy-induced senescence (TIS) is a state of permanent growth arrest and a powerful tumor suppressor mechanism elicited by several stressors, including chemotherapeutic agent^4,5^. In recent years, cancer immunotherapy with Immune Checkpoint Blockade (ICB) drugs has emerged as a revolutionary intervention for treating a variety of solid tumors by unleashing the cytotoxic potential of T cells against cancer cells^29^, although it had limited success in the context of most hematological malignancies. Here, we investigated the contribution of the interplay between cellular senescence and the adaptive immune system to the anti-tumor effects of conventional chemotherapy in longitudinally analyzed primary AML samples. By integrating gene expression analysis with cellular-based evaluation of senescent markers, we first identified two groups of AML patients based on their competence to accumulate SA-β-GAL to different extents upon *ex-vivo* chemotherapy and defined them as “Senescence High” or “Senescence Low” samples. Consistently, in “Senescence High” patients, we reported upregulation of HLA class I and II molecules and their regulators compared to untreated counterparts. We validated RNA-seq findings by measuring the expression levels of HLA molecules on cell surfaces on TIS AML primary blasts and found a positive correlation between senescence and HLA class I and II upregulation, suggesting that the two phenomena are interconnected. Conversely, AML samples from the “Senescence Low” group of patients did not display such upregulation, despite the similar effectiveness of chemotherapy in inducing proliferation arrest in both patient groups. One intriguing hypothesis is that these two groups of AML patients present distinct epigenetic profiles that may be responsible for the upregulation of HLA molecules class I and II observed upon senescence induction. Along this line, we recently reported in AML patients relapsing after allogeneic hematopoietic stem cell transplantation (allo-HSCT) epigenetic silencing of HLA class II molecules via the Polycomb Repressor Complex 2 (PRC2) and demonstrated that PRC2 inhibitor treatment can re-establish a proficient anti-leukemic T cell response^30^.

By undertaking several state-of-the-art immunological assays, we reported that “Senescence High” AML blasts undergoing TIS elicited a strong CD8^+^ T cell activation, likely due to HLA class I upregulation, and led to leukemia eradication in line with two recent reports showing that senescence triggers CD8^+^ T cells anti-tumor immunity^15,16^, supporting the idea that immune recognition of senescent cells by CD8^+^ T cells is conserved even in hematological cancers. HLA class II molecules play a central role in the control of adaptive immune responses through the activation of CD4^+^ T cells^31^. We discovered that HLA class II up-regulation in senescent AML blasts correlated with increased CD4^+^ T cell immune activation, as determined by enhanced proliferation, immunological synapsis formation, and eradication of TIS cells, an aspect not considered in studies performed so far in the context of immunogenicity of solid tumors^15,16^. The involvement of CD4^+^ T cells in AML eradication is in line with the reported role of this T cell subset in mediating immune recognition of post-allogeneic Hematopoietic Cells Transplant (allo-HCT) relapses upon reactivation of HLA class II expression^30^. Although the mechanisms are not fully understood, it would be interesting to evaluate whether the killing activity of CD4^+^ T cells against TIS cells may be related to the induction of apoptosis via FAS/FASL interaction, as reported in other settings^32^.

To date, whether senescent AML cells increase the presentation of tumor or senescence-specific antigens remains to be defined; however, based on the enhanced activation of T cells against senescent blasts compared to IFN-γ treatment, a well-established regulator of Class I and II molecules, one can speculate that TIS AML may present senescence-related antigens or coexpress on their surface costimulatory ligands that may further boost HLA-mediated T cell activation. Emerging evidence points to a link between senescent cells and epigenetic derepression of transposable elements including transcriptional reactivation of endogenous retroviruses (ERVs) ^33,34^ Interestingly, cancer cell treatment with DNA demethylating agents leads to the cytosolic accumulation of ERV nucleic acids activating the interferon pathway and boosting antitumor innate and adaptive immune responses^35,36^. Thus, aberrant accumulation of ERV nucleic acids or their presentation as viral antigens on HLA molecules may contribute to the increased immunogenicity of TIS AML via viral mimicry^37^.

In our study, we also reported that induction of senescence rather than genomic DNA damage *per se* is necessary to promote the upregulation of HLA class I and II molecules. Palbociclib, a Cyclin-Dependent Kinase (CDK) 4 and 6 inhibitor (CDK4/6i), can induce senescence and cell cycle arrest in the absence of physical DNA damage, working as a p16 mimetic^5^. We showed that CDK4/6i treatment in AML not only results in senescence-dependent upregulation of antigen processing and presentation but also translates into enhanced CD4^+^ and CD8^+^ T cell anti-leukemic immunity. Accordingly, in a mouse model of breast cancer, CDK4/6i treatment favored CD8^+^ T cell recruitment and increased cancer cell immunogenicity due to the reactivation and presentation of ERV proteins^38^. Since intensive chemotherapy may not be suitable for elderly patients, given the frequent presence of comorbidities, Palbociclib may represent a valid therapeutic alternative for AML patients, circumventing most side effects of high-dose chemotherapy.

While our data show that senescence induction shortly after chemotherapy renders AML cells highly immunogenic, there is also evidence that senescence may be detrimental in the long-term promoting leukemia stemness and chemoresistance^8^. Attempts to mitigate the long-term detrimental aspects of senescence in cancer have been the focus of intense scientific investigation^39^. Although their effects may be transient, selective SASP inhibitors may reduce senescence-related chronic inflammation that contributes to tumor growth and immune evasion. Instead, senolytic compounds exploit senescence vulnerabilities and have been shown to eliminate senescent cells in preclinical models^40^ and would be preferred to ameliorate chemotherapy-induced long-term side effects in cancer patients delaying or preventing relapse.

Nonetheless, their use in clinical studies has been challenged by their systemic organ and tissue toxicity^41^. Our study indicates that by potentiating the adaptive immune-system response against senescent cells, we may reach a higher level of senolysis, thus preventing the long-term detrimental impact of persistent senescent cells. In line with this, genetically-engineered CAR-T cells against a receptor upregulated on the surface of senescent cells ameliorated senescence-associated pathologies in mouse models^42^.

ICBs have successfully changed how we treat solid tumors although their efficacy in leukemia has been limited^29^. Here, we described an enhanced T cell activation for both T cell subsets when combining TIS and ICB compared to chemo or ICB monotherapy, suggesting that senescence competence may be exploited to stratify AML patients that will benefit from ICB treatment. Finally, given that allogeneic bone marrow transplantation for AML patients represents one of the first, and by large the most widely clinically implemented, demonstration of adoptive immunotherapy of cancer, it would be of high interest to address whether senescence competence upon chemotherapy may enhance the curative effects of allo-HCT.

In conclusion, our findings uncovered a novel link between therapy-induced senescence and leukemia immune recognition by the adaptive immune system, providing a rationale for senescence-based personalized immunotherapies for AML patients.

## METHODS

### Collection and purification of primary AML samples and *ex vivo* treatments

Primary AML patient samples were collected and cryopreserved at the San Raffaele Leukemia Biobank upon signing of a specific written informed consent, in accordance with the Declaration of Helsinki (Protocol “Banca Neoplasie Ematologiche” approved by the San Raffaele Ethic Committee on 10/05/2010, latest amendment on 06/14/12) and analyzed according to the “SEN-LEUK”Protocol (approved by the San Raffaele Ethic Committee on 12/10/2017). After thawing, cells were cultured on a layer of irradiated human fibroblasts (BJ-hTERT) in 5637 cell line-conditioned and cytokine-supplemented media. Blasts were purified by a CD34- or CD33-based magnetic-activated cell separation system (for CD34: cat. #130-046-702, for CD33: cat. #130-045-501, both from Miltenyi Biotec) after 3 days of pre-culturing or by FACS sorting using the following antibodies: Brilliant Violet 510™ hCD45 (cat#368526 from Biolegend), CD33-FITC (cat#IM1135U from Beckman Coulter), CD34-PE (cat#130-113-179 from Miltenyi Biotech). Following purification, AML cells were re-seeded and after 3 days media was refreshed and the stromal layer was replaced. ARA-C at 410nM (Stock solution: Citarabina Hos - Iniet 100mg/5ml, C.A.T. L01BC01 from Hospira) and CDK4/6i, also known as Palbociclib, at 10μM (cat#S1116 From Selleck Chemicals) were administered at times 0 and 72 hours. After 5 days of treatment (120 hours), fluorescence-based detection of SA-β-GAL was performed as described below.

### Senescence evaluation

Cells were seeded at 1×10^5^ cells/mL and incubated with Chloroquine (cat#C6628 from Sigma-Aldrich) at a final working concentration of 150 μM for 1h at 37°C. Then, 5-Dodecanoylaminofluorescein Di-β-D-Galactopyranoside (C_12_FDG, cat#D2893 from ThermoFisher) was added to a final working concentration of 16,66 nM for 30 min at 37°C. After washing in PBS, stained cells were analyzed by imaging-based flow cytometry technology (ImageStreamX) or by FACS at the CANTO II flow cytometer. To evaluate cell proliferation alongside senescence, following a 48-hour incubation with EdU (2 μM final concentration), cells were first stained for fluorescent-SA-β-GAL, then with Pacific Blue AnnexinV (cat#640918 from Biolegend) using Annexin binding buffer 1X (cat#556454 from BD). Cells were fixed, permeabilized and stained for EdU (Click-iT™ EdU Alexa Fluor™ 647 Flow Cytometry Assay Kit, cat#C10635 from ThermoFisher). After the staining, cells were analyzed as follows. Only events with a gradient value higher than 60 based on brightfield images were considered cells on focus. Single-cell events were selected by plotting the area against the aspect ratio. Alive cells (negative for AnnexinV) within the single-cell gate were selected for EdU signal and the senescence marker fluorescent-SA-β-GAL.

For classical, non-fluorescent, SA-β-GAL analyses, multitest slides (cat#096041505 from MP Biomedicals) were treated for 20 min RT with Poly-L-lysine solution 0.1 % (w/v) in H_2_O solution (cat# P8920 from Sigma-Aldrich). After two washes with PBS, approximately 1.5×10^5^ cells were seeded on covers for 20 min and fixed with a Paraformaldehyde solution 4% in PBS (cat#sc-281692 from Santa Cruz Biotechnology) for 20 min RT. Then X-GAL working solution (cat#9860S from Cell Signalling Technology) was added to the slide for 16 hours at 37°C. After two washes with PBS, nuclei were stained with DAPI (cat# D9564 from Sigma Aldrich), and covers were mounted with Aqua-Poly/Mount solution (cat#18606-5 from Polysciences Europe) on glass slides. Images were acquired using Widefields Zeiss Axio Observer.Z1 microscope.

### Total RNA seq library preparation and sequencing

Total RNA was isolated with miRNeasy Mini Kit (cat# 217004, from QIAGEN), according to manufacturer’s instructions. RNA was quantified with The Qubit 2.0 Fluorometer (ThermoFisher) and quality was assessed by Agilent 4200 TapeStation (Agilent Technologies). Minimum quality was defined as RNA integrity number (RIN) >7. Then, 10-30 ng of total RNA was used for library preparation with Illumina Stranded mRNA Prep Ligation (Illumina) and sequenced on a NextSeq 500 (Illumina).

### Bioinformatics and statistical analysis

High-quality data were obtained after reads quality inspection and adapter trimming with Trim Galore (v0,6). Reads were then aligned to the human reference genome (GENCODE version hg38 primary assembly) with STAR (v2.7.6a^43^). Gene expression quantification was obtained with featureCounts (v2.0.1^44^). The differential expression (DE) analysis was performed with the DESeq2 package^45^, testing for the contrast treatment vs control (ARA vs CTRL), and taking into account the paired design of the experiment (~ *treatment + patient*). Finally, adjusted *P* values for multiple hypothesis corrections were used to retain significant genes (FDR < 0.05). Over-Representation Analysis (ORA) of deregulated genes was performed with the function enrichGO from the clusterProfiler package^46^. Furthermore, Gene Set Enrichment Analysis (GSEA)^47^ was performed using the clusterProfiler package, ranking genes according to the Log_2_FC value. The analysis was performed considering the MSigDB database (Hallmark signatures lists, release 7.2) and literature available signatures, retaining results with FDR < 0.05. Figures were plotted with the ggplot2 package in R (v4.0.3). The analysis for the detection of HLA genes (class I and II) was performed by aligning high-quality reads to the human reference genome (GENCODE version GRCh38.p13, including patches and haplotypes) with a multi-mapping approach, considering the following parameters for the alignment with STAR ‘--winAnchorMultimapNmax 100 -- outFilterMultimapNmax 100’, and for the expression quantification with featureCounts ‘-M -- fraction’. The differential expression analysis was performed as described above.

### AML cell lines and senescence-inducing treatments

OCI-AML3 cells were maintained in Iscove’s Modification of DMEM (cat#10-016-CV from Corning) supplemented with 10% Fetal Bovine Serum, certified, One Shot™ format, United States (cat#A3160402 from Thermofisher), 1% Penicillin Streptomycin mixture (cat#DE17-602E from Lonza) and 2% L-Glutamine (cat#BE17-605E from Lonza) at 5% CO_2_ and 37°C. THP-1 cells (from ATCC) were maintained in RPMI 1640 (cat#10-040-CV from Corning) supplemented with 10% FBS, 2% P/S and 1% glutamine at 5% CO_2_ and 37°C. Treatments were performed by seeding cells in 6 wells plates either 1×10^5^ cells/mL with ARA-C 250nM (every other day for 7 days) or 3×10^5^ cells/mL with Palbociclib (CDK 4/6i) 1uM (every other day for 7 days); drug doses were chosen after conducting titration experiments. For growth curves, cells were seeded at 1×10^5^/ml in a 6 multiwell plate (3 technical replicates for each condition), then cells were counted with trypan blue solution using counting slides every other day. For viability assays, cells were FACS analyzed after live staining with PE Annexin V (cat#556422 from BD) combined with 7AAD (cat#420403 from Biolegend). This staining was performed using 1×10^5^ cells in 100μL of Annexin binding buffer 1X, adding 2 μL of PE Annexin V and 1μL of 7AAD. Stained cells were acquired at the CANTO flow cytometer, and data were analyzed using FlowJo software version 10.8.1 (Tree Star). Senescence evaluation was carried out as described above and HLA class I and II were quantified as described below.

### Cell cycle analysis

Cell cycle analysis was performed by combining 5-ethynyl-20-deoxyuridine (EdU) and Hoechst staining. The staining was performed with Click-iT Plus EdU 647 Flow Cytometry Assay Kit (supplied with all the reagents named below) in combination with Hoechst 33342. Practically, 2 μM EdU was added to the culture media of 3-5×10^4^ cells for 4 hours. Cells were washed in FACS buffer, centrifuged at 500 g for 5 min, and resuspended in 100 μl of 4% paraformaldehyde (PFA) for fixation. After 15 min at RT samples were washed and permeabilized with Saponin 15 min at RT. According to the manufacturer’s instructions, 500 μl of Click-iT Plus reaction cocktail was added to the samples and incubated for 30 min at RT. Finally, DNA was labeled with 2,5 μM Hoechst 33342 Solution overnight at 4°C protected from light. Sample acquisition was performed on FACSymphony A5 SORP and the collected data were analyzed using FlowJo software version 10.8.1 (Tree Star).

### HLA class I and II quantification

For immunophenotypic analysis, a maximum of 0.5×10^5^ cells per tube were stained in 100 μl of 1X PBS, 2% FBS, plus the mix of antibodies. Staining was performed at RT for 15 minutes, followed by washing with 2 ml of 1X PBS, 2% FBS before the analysis. Here, is the complete list of antibodies (all from Biolegend): HLA-ABC Pacific Blue (Clone W6/32, cat#311418), HLA-DR FITC (Clone L243, cat#307604) or HLA-DR APC-Cy7 (cat#307618) when combined to Senescence evaluation or T-cells assays. Signal was acquired using the FACS Aria Canto II flow cytometer (BD Biosciences). Data were processed using FlowJo software version 10.8.1 (Tree Star).

### RT-qPCR quantification of gene transcripts

Total RNA was extracted from live cells with miRNeasy Mini Kit according to the manufacturer’s instructions. 500 ng of total RNA (as quantified with Nanodrop 8000 device (Thermo Scientific)) was used for retrotranscription using iScript cDNA Synthesis Kit (cat#170-8891 from Bio-Rad) according to manufacturer’s instructions. An estimated 25ng of cDNA in 20μl reaction volume was analyzed in triplicate by quantitative RT-PCR amplification on a Viia7 Real-time PCR applied biosystem (Life Technology) using SYBR Green assay (QuantiFast SYBR Green PCR Kit, cat#4385618 from ThermoFisher) according to manufacturer’s instructions and for 40 cycles. CT-values were obtained by calculating the second derivative using Quant studios RT-PCR software and normalizing samples against a housekeeping gene *GUSB*. Following forward and reverse primers were designed with Metabion online software against *Homo Sapiens*. A primer efficiency analysis was performed to define the correct concentration according to the Leivac distribution.

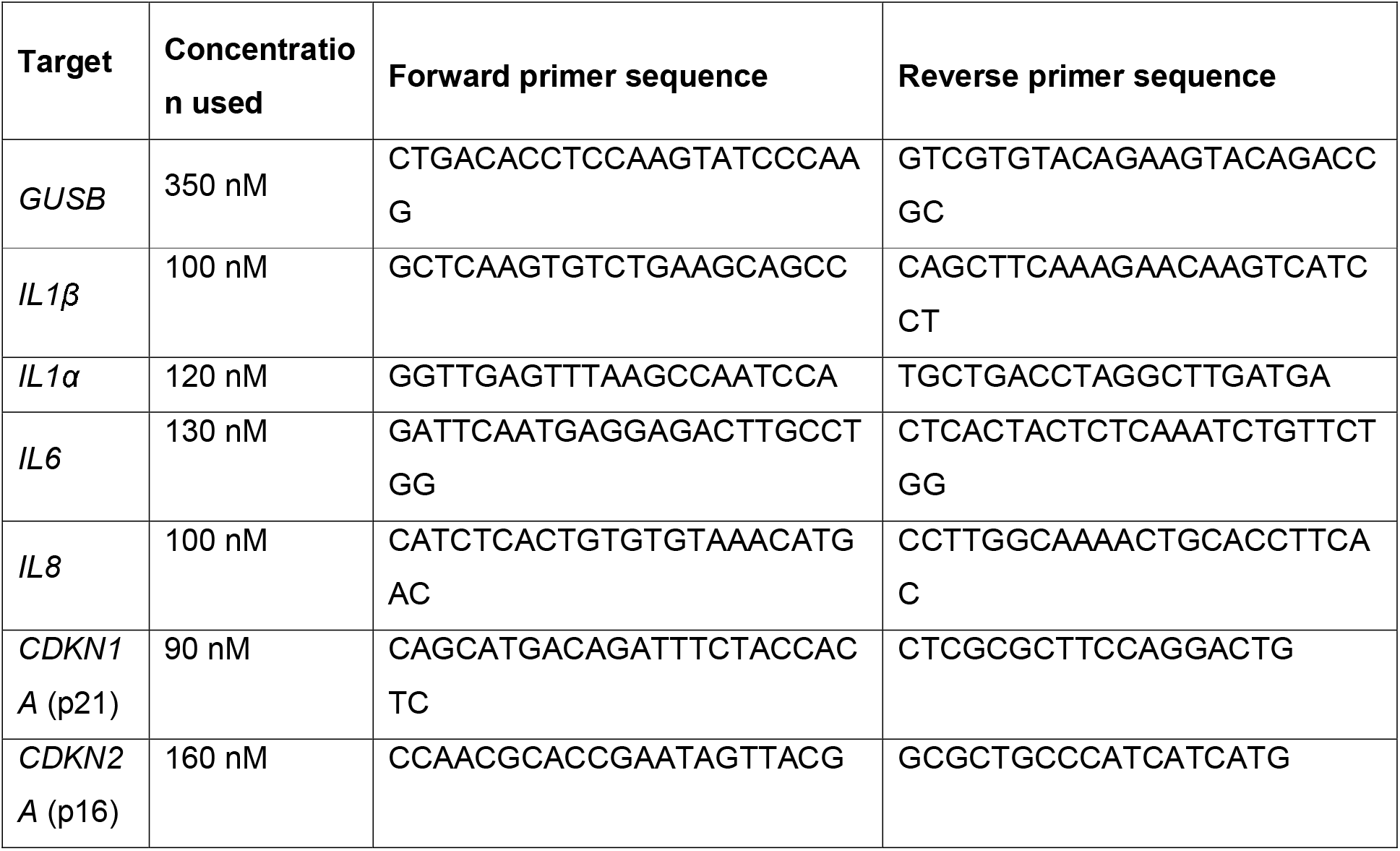

### Western blotting

Samples were loaded on 12 % Acrylamide gels. After protein fractionation by SDS-page samples were transferred to a nitrocellulose membrane (cat#10600002 from GE healthcare) and incubated with primary antibody solutions overnight at 4 °C (p21, mouse, cat#556430 from BD; p53, mouse, cat#SC-126 from Santa Cruz Biotechnology; Phospho-p53 (Ser15), rabbit, cat#9284S from Cell Signalling Technology; alpha Tubulin, rabbit, cat#ab4074 from Abcam; 17hosphor-Histone H2A.X (Ser139), mouse, cat#05-636 from Merck Millipore; Histone H3, rabbit, cat#ab1791 from Abcam) in PBS supplemented with 0,1 % Tween-20 (cat# P1379 from Sigma Aldrich) and 2% BSA (Cat#2153 from Sigma Aldrich). Membranes were washed three times with PBS-Tween 0.1 % for 10 min, then incubation with HRP-linked secondary antibody solution was carried on for 1 hour (according to primary antibody host species: Goat anti-Rabbit IgG (H+L) Secondary Antibody, HRP, cat#65-612-0 or Goat anti-Mouse IgG (H+L) Secondary Antibody, HRP, cat#62-652-0, both from Thermofisher). After 3 washes with PBS-Tween 0.1 % for 10 min, HRP substrate was applied (Lumi-Light Western Blotting Substrate cat#12015200001 from Sigma Aldrich, Immobilon Crescendo cat#WBLUR0020 from Merck Millipore, SuperSignal West Femto Maximum Sensitivity Substrate cat#34096 from ThermoFisher) to detect signal. Protein size was identified by matching bands from a colored protein ladder (cat#1610374 from Bio-Rad).

### Immunofluorescence analysis

Multitest slides were treated for 20 min with Poly-L-Lysine solution RT. After two washes with PBS, approximately 1.5×10^5^ cells were seeded on covers for 20 min and fixed with 4% paraformaldehyde for 20 min RT. Next, cells were permeabilized with 0.2% Triton X100 and probed with the indicated primary antibodies (Histone H2A.X (Ser139), mouse, cat#05-636 from Merck Millipore; 53BP1, rabbit, cat# A300-272A from Bethyl and Phospho-ATM S1981, rabbit, cat#2851S from Cell Signalling Technology) after blocking with 0.5% BSA and 0.2% Gelatin from cold water fish skin (cat#G7765 from Sigma Aldrich) in PBS. After primary antibodies incubation, we performed three washes with PBS and incubated with Alexa 488-, 568- and/or 647-labeled secondary antibodies (Donkey anti-Mouse IgG (H+L) Highly Cross-Adsorbed Secondary Antibody, Alexa Fluor™ 488, cat#A21202; Donkey anti-Rabbit IgG (H+L) Highly Cross Adsorbed Secondary Antibody, Alexa Fluor 647, cat#A31571 and Donkey anti-Rabbit IgG (H+L) Highly Cross-Adsorbed Secondary Antibody, Alexa Fluor 488, cat#A21206; all from Thermofisher). Nuclear DNA was stained with DAPI and covers were mounted with Aqua-Poly/Mount solution on glass slides. Fluorescent images were acquired using Widefields Zeiss Axio Observer.Z1 microscope or at SP5 Confocal.

### T cells purification

Whole peripheral blood samples from either healthy donors or AML patients were processed to obtain mononuclear cells (PBMCs) through density gradient centrifugation using Lympholyte™ (cat#DVCL5020-10 from Lonza). CD3^+^ T lymphocytes were isolated via magnetic separation with a Pan T cell Isolation Kit (cat#130-096-535 from Miltenyi Biotech) according to the manufacturer’s instructions. The negative fraction was collected. After the selection, viable samples consisted of >90% CD3^+^ cells. To discriminate T cell subsets (CD4^+^ and CD8^+^) we used specific antibodies listed below.

### Mixed Lymphocyte Culture and downstream analyses

Purified T cells were stimulated with leukemic blasts from cell lines or primary samples at an effector:target ratio of 1:1. When indicated by the experimental setting, blasts were pre-treated for 48 hours with Recombinant Human IFN-γ at a final concentration of 200 U/mL (cat#300-02 from PeproTech). Cells were cultured in IMDM supplemented with 1% L-glutamine (G), 1% penicillin/streptomycin (P/S), 10% Human Serum (cat#ECS0219D from Euroclone) and IL-2 (cat#F027131010 from Novartis) at a final concentration of 150 UI/mL. IL-2 was replaced every 3–4 days. After stimulation, T cells were tested against targets of choice co-culturing them at a 1:1 ratio. For the HLA-DR blockade condition cells were pre-treated with 10 μg/mL HLA-II blocking antibody (Bio X Cell, Lebanon, NH, BE0306) or 10 μg/ml isotype IgG antibody in the absence of FBS overnight at 37°C. For the ICB conditions, at the start of the co-culture cells were treated with 10 μg/mL Ipilimumab (anti-CTLA-4, cat#A2001 from Selleck Chemicals), Nivolumab (anti-PD-1, cat#A2002 from Selleck Chemicals) or Durvalumab (anti-PD-L1, cat# A2013 from Selleck Chemicals).

T-cells were then collected at different time points according to downstream analyses: proliferation, degranulation, killing activity and immunological synapse formation. For T cell proliferation, before co-culture T cells were resuspended 1×10^6^ cells/mL in 1X PBS containing Celltrace Violet Cell (cat#C34557 from Thermofisher) for 20 minutes at 37°C, then washed with pre-heated supplemented IMDM supplemented. Dilution of Cell Trace Violet was quantified by FACS analyses 5 days after co-culture using primed T cells cultured alone as a negative control.

T-cell degranulation was evaluated by adding an anti-CD107a antibody at the beginning of coculture (CD107a-FITC clone H4A3, BD, Catalog N° 555800) and Golgi Stop (BD, Catalog N° 554724) after 3 hours. At 6 hours after co-culture cells were stained with CD8-PECy7 and CD4-PE antibodies, then the signal was acquired at the FACS Canto II.

For the evaluation of AML blasts-killing, before co-culture blasts were stained for senescence evaluation as described above and then mixed to T cells in a ratio 1:1 in a 6 multiwell size plate (cell total number 1×10^6^ cells/mL). After the co-culture start, IncuCyte caspase-3/7 Red (cat#4704 from Sartorius) was added in the media. The plate was then inserted inside IncuCyte (Sartorius) instrument and the co-culture was followed for 3 days acquiring images every hour. To discriminate T cell killing by cell-autonomous induction of apoptosis in AML, a control condition of AML blasts alone was analyzed.

For the evaluation of immunological synapses formation, 4 hours after co-culture, C_12_FDG staining was performed. Cells were then stained with anti-CD3 and anti-CD33 human antibodies, fixed in PFA 2% and stained with phalloidin conjugated with AlexaFluor647 (cat# A22287, Thermo Scientific). To find the contact zone between blasts and T cells, total events were gated on true Blast/ T-cell pairs (doublets positive for both CD33 and CD3 signals). The area of the immunological synapse was then identified in the two-cell conjugates using two independent algorithmic functions. The Valley function identifies the dimmest area between two nuclei and the Interface function identifies the point of contact between two cells and selects only the (CD3^+^) T cell portion of the T cell/blast synapse^48^. Data acquisition (1×10^5^ cells per sample) was performed with the ImageStreamX system. Single color controls were collected and used to calculate a spectral crosstalk compensation matrix to compensate for the images, and a mask to detect the enrichment of actin signal in the interface of two cells was designed by using IDEAS 3.0 software.

### Characterization of T cell exhaustion and activation markers

For immunophenotypic analysis, a maximum of 0.5×10^5^ cells per tube were stained in 100 μl of 1X PBS, 2% FBS, plus the mix of antibodies. Staining was performed at RT for 15 minutes, followed by washing with 2 mL of 1X PBS and 2% FBS before the analysis. Here, is the complete list of antibodies used for experiments of T cell activation or exhaustion against primary AML: Anti-CD279 (PD-1) FITC (cat#329904 from Biolegend), CD366 (TIM-3) BV421 (cat#345008 from BioLegend), CD223 (LAG-3) BV605 (cat#369324 from BioLegend), CD3 BUV737 (cat#565466 from BD), CD4 BUV395 (cat#563550 from BD), CD8 APCH7 (cat#641400 from BD), CD69 APC (cat#310910 from BioLegend), CD25 PE (cat#130-113-286 from Miltenyi Biotec). Signal was acquired using FACSymphony™ flow cytometer (BD Bioscience). Data were processed using FlowJo version 10.8.1 (Tree Star) and SPICE^27^.

Here, is the complete list of antibodies used for experiments of co-culture between T cells and AML cell lines: CD8 PE-Cy7 (cat#557746 from BD), CD4 FITC (cat#980802 from BioLegend) or CD4 PE (cat#300508 from BioLegend) when combined to T-cells degranulation assay, CD69 APC (cat#310910 from BioLegend), CD25 PE (cat#130-113-286 from Miltenyi Biotec), ZOMBIE Aqua (cat#423102 from BioLegend). Signal was acquired using the FACS Aria Canto II flow cytometer (BD Biosciences). Data were processed using FlowJo software version 10.8.1 (Tree Star).

## Acknowledgments

The Authors would like to thank all members of Di Micco’s laboratory for discussion; Martina Ciccimarra and Laura Zito for technical suggestions on Myxed Lymphocyte Culture assays, the FRACTAL and ALEMBIC facilities at San Raffaele Hospital. D.G and S.F. conducted this study as partial fulfillment of their Ph.D. in Molecular Medicine, International Ph.D. School, Vita-Salute San Raffaele University, Milan, Italy. A.C. is a European Hematology Association and American Society of Hematology Translational Research Training in hematology (EHA and ASH-TRTH) early career hematological scientist. A.S. is funded by EMBO Postdoctoral Fellowship (awarded number ALTF 245-2022). Research in the R.D.M laboratory is supported by the Associazione Italiana per la Ricerca sul Cancro (My first AIRC Grant MFAG 2019 –PI ID.23321), by a Career Development Award from Human Frontier Science Program (HFSP) by an Advanced Research Grant from the European Hematology Association (EHA) by a Hollis Brownstein Research Grant from Leukemia Research Foundation (LRF), by the New York Stem Cell Foundation and the European Research Council (consolidator grant 101003186, ReviveSTEM). R.D.M is a New York Stem Cell Foundation Robertson Investigator.

## Author contributions

**D. Gilioli:** Conceptualization, data curation, formal analysis, investigation, methodology, writing–original draft, writing–review and editing. **S. Fusco:** Data curation, formal analysis, investigation, methodology, writing–original draft, writing–review and editing. **T. Tavella:** Data curation, formal analysis, methodology, writing–review and editing. **K. Giannetti:** Data curation, formal analysis, methodology, writing–review and editing. **A. Conti:** Data curation, formal analysis, methodology. **A. Santoro:** Data curation, formal analysis, methodology. **E. Carsana:** Formal analysis, methodology, **S. Beretta:** Methodology. **M. Schönlein:** Data curation, formal analysis, methodology. **V. Gambacorta:** methodology. **F. Aletti**: Data curation, Resources. **M. Carrabba**: Data curation, Resources. **C. Bonini**: Methodology. **F. Ciceri:** Resources. **I. Merelli:** Data curation, formal analysis, supervision, methodology, writing–review and editing. **L. Vago:** Methodology, Data curation, writing–review and editing. **C. Schmitt:** Methodology, Data curation, writing–review and editing. **R. Di Micco:** Conceptualization, resources, supervision, funding acquisition, data curation, investigation, methodology, writing–original draft, project administration, writing–review and editing.

## SUPPLEMENTARY FIGURE LEGENDS

**Supplementary Figure S1. Flow cytometry gating strategies and principal component analysis of RNA sequencing in primary AML samples**. **A,** Gating strategy and representative flow cytometry plots with the sequential logical gates of primary AML blasts purified by either FACS or a magnetic-activated cell separation system sorting after 3 days of pre-culturing. At first, “Single cells” were identified, then “Alive cells” were gated according to physical parameters. Based on the expression of human CD45 and SSC-A, blasts were identified within the CD45dim cell population (“Blasts”) and further distinguished by the expression of CD33 or CD34. When double positive, magnetic separation was carried out through CD34-based magnetic-activated cell separation, otherwise CD33-based magnetic-activated cell separation was applied. **B,** Principal Component Analysis (PCA) of regularized log2 counts after batch correction by limma of 6 samples in paired design and color-coded by treatment control (CTRL, red), ARA-C (ARA-C, blue) from the RNA-seq data obtained from 3 patients: UPN03, UPN06 and UPN09.

**Supplementary Figure S2. Senescence establishment in primary AML and cell lines. A,** Schematic representation of cell line treatment upon ARA-C (250nM) administration every other day. **B-C,** Growth curve of THP-1 (B) and OCI-AML3 (C) cells either untreated (CTRL, red) or treated (ARA-C, blue). 100.000 cells were seeded at the starting point of the growth curve (n=5). **D-E,** Vitality assay of untreated (CTRL, red) and treated (ARA-C, blue) THP-1 (D) and OCI-AML3 (E) cells by AnnexinV/7AAD staining. Curves represent the percentage of alive (double negative) cells over time. **F**, Western Blot analysis for p53, phospho-p53 (Ser15) and p21 in untreated (CTRL) or chemotherapy-treated (ARA-C) OCI-AML3 (on the left) and THP-1 (on the right) cells. **G-H**, Gene expression analysis of canonical SASP genes (*IL8, IL6, CCL2, IL1β* and *IL1α*) in CTRL (red) or ARA-C treated (blue) THP-1 (G) and OCI-AML3 (H) cells. **I-J**, Representative images of SA-β-GAL in untreated (CTRL, top row) or treated (ARA-C, bottom row) in THP-1 (I) and OCI-AML3 (J) cells. **K**, Quantification of SA-β-GAL positive cells from panel I-J. **L**, Representative confocal images of, from left to right, DAPI, γH2AX and 53BP1 staining in primary AML blasts obtained from UPN21 at diagnosis (Day 0) or 4 days after *in-vivo* chemotherapy start (Day 4). **M**, Representative confocal images of phospho-ATM (S1981) staining in primary AML blasts obtained from UPN21 at diagnosis (Day 0) or 4 days after *in-vivo* chemotherapy start (Day 4).

**Supplementary Figure S3. Immune-related categories and HLA-class I or HLA-class II levels upon senescence induction in primary AML. A**, Over-Representation Analysis (ORA) of positive Differentially Expressed Genes (DEGs). Dot plot of the GO terms reporting selected enriched terms for the Molecular Function (MF) and Cellular Component (CC) ontology branches. **B**, Distribution of cells along the fluorescent intensity axis corresponding to HLA-DR (on the left) or HLA-ABC (on the right) expression. Two representative images with results from Senescence High and Senescence Low patients either unstained (grey), CTRL (red) or ARA-C treated (blue) are shown.

**Supplementary Figure S4. Chronic or acute effects of senescence-inducing treatments in AML cell lines. A,** Schematic representation of THP-1 cells treatment, each arrow represents a single ARA-C (250 nM) or CDK 4/6i (1 μM) administration. **B,** Cell cycle evaluation of THP-1 (left) and OCI-AML3 (right) cells untreated (CTRL) or treated with ARA-C or CDK 4/6i after 3 days from seeding. Cell cycle phases (G2 in light violet, S in violet and G1 in dark violet) were evaluated by combining EdU with Hoechst staining. **C-D,** Growth curve of THP-1 (C) and OCI-AML3 (D) cells either untreated (CTRL, red) or treated with ARA-C (blue) or CDK 4/6i (green). 100.000 cells were seeded at the starting point of the growth curve (n=5). **E-F**, Percentage of alive cells in THP-1 (E) and OCI-AML3 (F) cells either untreated (CTRL, red) or treated with ARA-C (blue) or CDK 4/6i (green) 48 and 168 hours after seeding. **G**, Western Blot analysis of γH2AX in control (CTRL), ARA-C or CDK 4/6i treated cells. Histone H3 was used as loading control. **H**, On the left, distribution of cells along the fluorescent intensity axis corresponding to SA-β-GAL activity upon C_12_FDG staining (see Methods section). Here, a representative image with results from samples either untreated (CTRL, red) or treated with either ARA-C (blue) or CDK 4/6i (green) are shown. On the right, percentage of fluorescent SA-β-GAL positive cells from untreated (CTRL, red) and ARA-C (blue) or CDK 4/6i (green) treated AML cells are shown for each group. Individual dots represent biological replicates. Statistical test: Kruskal-Wallis with Dunn’s multiple comparison. ‘**’p < 0.005; ‘****’p < 0.0001. **I**, On the left, distribution of cells along the fluorescent intensity axis corresponding to HLA-ABC expression. Representative images with results from untreated (CTRL, red) and ARA-C (blue) or CDK 4/6i (green) treated AML cells are shown for each group. On the right, the percentage of HLA-ABC positive cells from untreated (CTRL, red) and ARA-C (blue) or CDK 4/6i (green) treated AML cells are shown for each group. Individual dots indicate biological replicates. **J**, On the left, representative images of FACS plots showing HLA-DR signal against SSC from AML cells either untreated (CTRL, red) or treated with either ARA-C (blue) or CDK 4/6i (green). On the right, the percentage of HLA-DR positive cells from untreated (CTRL, red) and ARA-C (blue) or CDK 4/6i (green) treated AML cells are shown for each group. Individual dots indicate biological replicates. Statistical test: Kruskal-Wallis with Dunn’s multiple comparisons. ‘*’p < 0.05; ‘**’p < 0.005. **K-L,** Percentage of fluorescent SA-β-GAL (C_12_FDG, J) or HLA-DR (L) positive cells 24 hours upon treatment in untreated (CTRL, in red) or ARA-C (blue) or CDK 4/6i (green) treated cells. Individual dots represent biological replicates. **M**, Mean Fluorescent Intensity (MFI) of HLA class I positive cells 24 hours upon treatment in untreated (CTRL, in red) or ARA-C (blue) or CDK 4/6i (green) treated cells.

**Supplementary Figure S5. Analyses of senescence-induced T cell activation. A**, Gating strategy and representative flow cytometry plots with the sequential logical gates of T cells. At first, “Single cells” were identified, then “Alive cells” were gated according to physical parameters and Zombie (BV510) staining (negative cells are identified as alive). Different T cell subsets were identified as CD4^+^ (FITC) or CD8^+^ (PE-Cy7) when positive for only one marker. **B,** Gating strategy used for T cell proliferation evaluation. Representative flow cytometry plots with the sequential logical gates to identify CD8^+^ and CD4^+^ cells are shown in panel A. Then, according to CT-Violet (PB) signal and using an unstained sample, the CTV negative (CTV^-^) population was identified. Here, representative flow cytometry plots from samples of the co-culture are shown, from left to right: T cells ALONE, T cells + CTRL cells, T cells + SEN by ARA-C cells, T cells + SEN by ARA-C cells with HLA-DR blockade. **C-D**, Gating strategy used for T cell activation evaluation. Representative flow cytometry plots with the sequential logical gates to identify CD8^+^ and CD4^+^cells are shown in panel A. Then, CD69 (C) and CD25 (D) positive cells were identified according to, respectively, APC or PE signal. Here representative images with results from samples of the MLR assay are shown, T cells + CTRL cells (left), T cells + SEN by ARA-C cells (right), **E,** Gating strategy used for T cell degranulation evaluation. Representative flow cytometry plots with the sequential logical gates to identify CD8^+^ and CD4^+^ cells are shown in panel A. Then, CD107a (FITC) positive cells were identified. Here, representative images with results from samples of the co-culture assay are shown, from left to right: T cells alone, T cells + CTRL cells, T cells + SEN cells. **F,** Gating strategy for immunological synapses evaluation on ImageStreamX. On the upper row, the gating strategy to evaluate immunological synapses is shown, while on the bottom row, total T-cells are shown. Upper Row: Doublets were identified by selecting events with “Area on BF signal” higher than 60 (taking only events in focus) and “Aspect ratio” lower than 0.8 (aspect ratio is used to evaluate the circularity of an event: if the value is near 1, the event is circular; below 0,6 it is asymmetric). Within doublets, double positive events were used to identify contacts between T cells (CD3) and AML blasts (CD33). In this population, the number of immunological synapses was identified based on a higher actin signal. Bottom Row: Whole cells were identified by selecting events with “Area on BF signal” higher than 60 (taking only events in focus) and “Aspect ratio” lower than 1. Then, total T-cells were identified as CD3^+^.

**Supplementary Figure S6. Activation and exhaustion profiles of T cells from AML patients at diagnosis or 30 days post-chemotherapy. A-B**, Percentage of CD69^+^ (A) and CD25^+^ (B) T cells collected at diagnosis (Day 0) and 30 days after chemotherapy start (Day 30) were analyzed in samples from AML patients belonging to either the Senescence High (UPN10, UPN11, UPN13, UPN14) or the Senescence Low (UPN12) groups. **C**, T cell exhaustion markers (PD-1, LAG-3, TIM-3) in AML patients from the Senescence High (UPN10, UPN11, UPN13, UPN14) or Senescence Low (UPN12) groups. The pie chart represents the percentage of T cells positive for 3 (red), 2 (orange), 1 (yellow) or 0 (green) inhibitory markers. Arcs indicate the percentage of cells positive for the specific marker indicated: PD-1 (light blue), LAG-3 (blue) and TIM-3 (purple). Charts were obtained using SPICE software.

**Supplementary Table S1. Characteristics of AML patients considered in the study.**

**Supplementary Table S2. Table with DEGs and GSEA analyses from RNA-sequencing dataset.**

**Supplementary Table S3. Table with GO, KEGG and HLA analyses from RNA-sequencing dataset.**

## Notes

### Competing Interest Statement

The authors have declared no competing interest.

## REFERENCES

1. Wiernik, P. H. & Dutcher, J. P. Clinical importance of anthracyclines in the treatment of acute myeloid leukemia. Leukemia 6 Suppl 1, 67–69 (1992).

2. Giordano, T. J. The cancer genome atlas research network: a sight to behold. Endocr Pathol 25, 362–365 (2014).

3. di Micco, R., Krizhanovsky, V., Baker, D. & d’Adda di Fagagna, F. Cellular senescence in ageing: from mechanisms to therapeutic opportunities. Nat Rev Mol Cell Biol 22, 75–95 (2021).

4. Lee, S. & Schmitt, C. A. The dynamic nature of senescence in cancer. Nat Cell Biol 21, 94–101 (2019).

5. Wang, L., Lankhorst, L. & Bernards, R. Exploiting senescence for the treatment of cancer. Nat Rev Cancer 22, 340–355 (2022).

6. Ewald, J. A., Desotelle, J. A., Wilding, G. & Jarrard, D. F. Therapy-induced senescence in cancer. J Natl Cancer Inst 102, 1536–1546 (2010).

7. Demaria, M. et al. Cellular Senescence Promotes Adverse Effects of Chemotherapy and Cancer Relapse. Cancer Discov 7, 165–176 (2017).

8. Duy, C. et al. Chemotherapy Induces Senescence-Like Resilient Cells Capable of Initiating AML Recurrence. Cancer Discov 11, 1542–1561 (2021).

9. Milanovic, M. et al. Senescence-associated reprogramming promotes cancer stemness. Nature 553, 96–100 (2018).

10. Burton, D. G. A. & Stolzing, A. Cellular senescence: Immunosurveillance and future immunotherapy. Ageing Res Rev 43, 17–25 (2018).

11. Mevorach, D. et al. What do we mean when we write ‘senescence,’ ‘apoptosis,’ ‘necrosis,’ or ‘clearance of dying cells’? Ann N Y Acad Sci 1209, 1–9 (2010).

12. Egashira, M. et al. F4/80+ Macrophages Contribute to Clearance of Senescent Cells in the Mouse Postpartum Uterus. Endocrinology 158, 2344–2353 (2017).

13. Soriani, A. et al. ATM-ATR-dependent up-regulation of DNAM-1 and NKG2D ligands on multiple myeloma cells by therapeutic agents results in enhanced NK-cell susceptibility and is associated with a senescent phenotype. Blood 113, 3503–3511 (2009).

14. Arora, S. et al. Invariant Natural Killer T cells coordinate removal of senescent cells. Med (N Y) 2, 938–950.e8 (2021).

15. Chen, H.-A. et al. Senescence rewires microenvironment sensing to facilitate anti-tumor immunity. Cancer Discov (2022) doi:10.1158/2159-8290.CD-22-0528.

16. Marin, I. et al. Cellular senescence is immunogenic and promotes anti-tumor immunity. Cancer Discov (2022) doi:10.1158/2159-8290.CD-22-0523/710086/CELLULAR-SENESCENCE-IS-IMMUNOGENIC-AND-PROMOTES.

17. Kang, T. W. et al. Senescence surveillance of pre-malignant hepatocytes limits liver cancer development. Nature 479, 547–551 (2011).

18. Jochems, F. et al. The Cancer SENESCopedia: A delineation of cancer cell senescence. Cell Rep 36, (2021).

19. Saul, D. et al. A new gene set identifies senescent cells and predicts senescence-associated pathways across tissues. Nat Commun 13, (2022).

20. Lee, S. et al. Virus-induced senescence is a driver and therapeutic target in COVID-19. Nature 599, 283–289 (2021).

21. Pribluda, A. et al. A senescence-inflammatory switch from cancer-inhibitory to cancer-promoting mechanism. Cancer Cell 24, 242–256 (2013).

22. Coppé, J. P., Desprez, P. Y., Krtolica, A. & Campisi, J. The senescence-associated secretory phenotype: the dark side of tumor suppression. Annu Rev Pathol 5, 99–118 (2010).

23. Purcell, M., Kruger, A. & Tainsky, M. A. Gene expression profiling of replicative and induced senescence. Cell Cycle 13, 3927–3937 (2014).

24. Fridman, A. L. & Tainsky, M. A. Critical pathways in cellular senescence and immortalization revealed by gene expression profiling. Oncogene 27, 5975–5987 (2008).

25. Biran, A. et al. Quantitative identification of senescent cells in aging and disease. Aging Cell 16, 661–671 (2017).

26. Toffalori, C. et al. Immune signature drives leukemia escape and relapse after hematopoietic cell transplantation. Nat Med 25, 603–611 (2019).

27. Roederer, M., Nozzi, J. L. & Nason, M. C. SPICE: exploration and analysis of post-cytometric complex multivariate datasets. Cytometry A 79, 167–174 (2011).

28. Salik, B., Smyth, M. J. & Nakamura, K. Targeting immune checkpoints in hematological malignancies. Journal of Hematology & Oncology 2020 13:1 13, 1–19 (2020).

29. Korman, A. J., Garrett-Thomson, S. C. & Lonberg, N. The foundations of immune checkpoint blockade and the ipilimumab approval decennial. Nature Reviews Drug Discovery 2021 21:7 21, 509–528 (2021).

30. Gambacorta, V. et al. Integrated Multiomic Profiling Identifies the Epigenetic Regulator PRC2 as a Therapeutic Target to Counteract Leukemia Immune Escape and Relapse. Cancer Discov 12, 1449–1461 (2022).

31. Oh, D. Y. & Fong, L. Cytotoxic CD4+ T cells in cancer: Expanding the immune effector toolbox. Immunity 54, 2701–2711 (2021).

32. Malyshkina, A. et al. Fas Ligand-mediated cytotoxicity of CD4+ T cells during chronic retrovirus infection. Sci Rep 7, (2017).

33. De Cecco, L., Dugo, M., Canevari, S., Daidone, M. G. & Callari, M. Measuring microRNA expression levels in oncology: from samples to data analysis. Crit Rev Oncog 18, 273–287 (2013).

34. de Cecco, M. et al. L1 drives IFN in senescent cells and promotes age-associated inflammation. Nature 566, 73–78 (2019).

35. Chiappinelli, K. Inhibiting DNA methylation causes an interferon response in cancer via dsRNA including endogenous retroviruses. Cell 162, 974–986 (2015).

36. Roulois, D. DNA-demethylating agents target colorectal cancer cells by inducing viral mimicry by endogenous transcripts. Cell 162, 961–973 (2015).

37. Jansz, N. & Faulkner, G. J. Endogenous retroviruses in the origins and treatment of cancer. Genome Biology 2021 22:1 22, 1–22 (2021).

38. Goel, S. et al. CDK4/6 inhibition triggers anti-tumour immunity. Nature 548, 471–475 (2017).

39. Chaib, S., Tchkonia, T. & Kirkland, J. L. Cellular senescence and senolytics: the path to the clinic. Nature Medicine 2022 28:8 28, 1556–1568 (2022).

40. Chang, J. et al. Clearance of senescent cells by ABT263 rejuvenates aged hematopoietic stem cells in mice. Nat Med 22, 78–83 (2016).

41. Grosse, L. et al. Defined p16High Senescent Cell Types Are Indispensable for Mouse Healthspan. Cell Metab 32, 87–99.e6 (2020).

42. Amor, C. et al. Senolytic CAR T cells reverse senescence-associated pathologies. Nature 583, 127–132 (2020).

43. Dobin, A. et al. STAR: ultrafast universal RNA-seq aligner. Bioinformatics 29, 15–21 (2013).

44. Liao, Y., Smyth, G. K. & Shi, W. featureCounts: an efficient general purpose program for assigning sequence reads to genomic features. Bioinformatics 30, 923–930 (2014).

45. Love, M. I., Huber, W. & Anders, S. Moderated estimation of fold change and dispersion for RNA-seq data with DESeq2. Genome Biol 15, (2014).

46. Wu, T. et al. clusterProfiler 4.0: A universal enrichment tool for interpreting omics data. Innovation (Cambridge (Mass.)) 2, (2021).

47. Subramanian, A. et al. Gene set enrichment analysis: a knowledge-based approach for interpreting genome-wide expression profiles. Proc Natl Acad Sci U S A 102, 15545–15550 (2005).

48. Ahmed, F., Friend, S., George, T. C., Barteneva, N. & Lieberman, J. Numbers matter: quantitative and dynamic analysis of the formation of an immunological synapse using imaging flow cytometry. J Immunol Methods 347, 79–86 (2009).

