## Supplementary figures and images for "Therapy-induced senescence upregulates antigen presentation machinery and triggers anti-tumor immunity in Acute Myeloid Leukemia"

### Supplementary figure 1

Fig. S1 Gating strategies and principal component analysis of primary AML samples

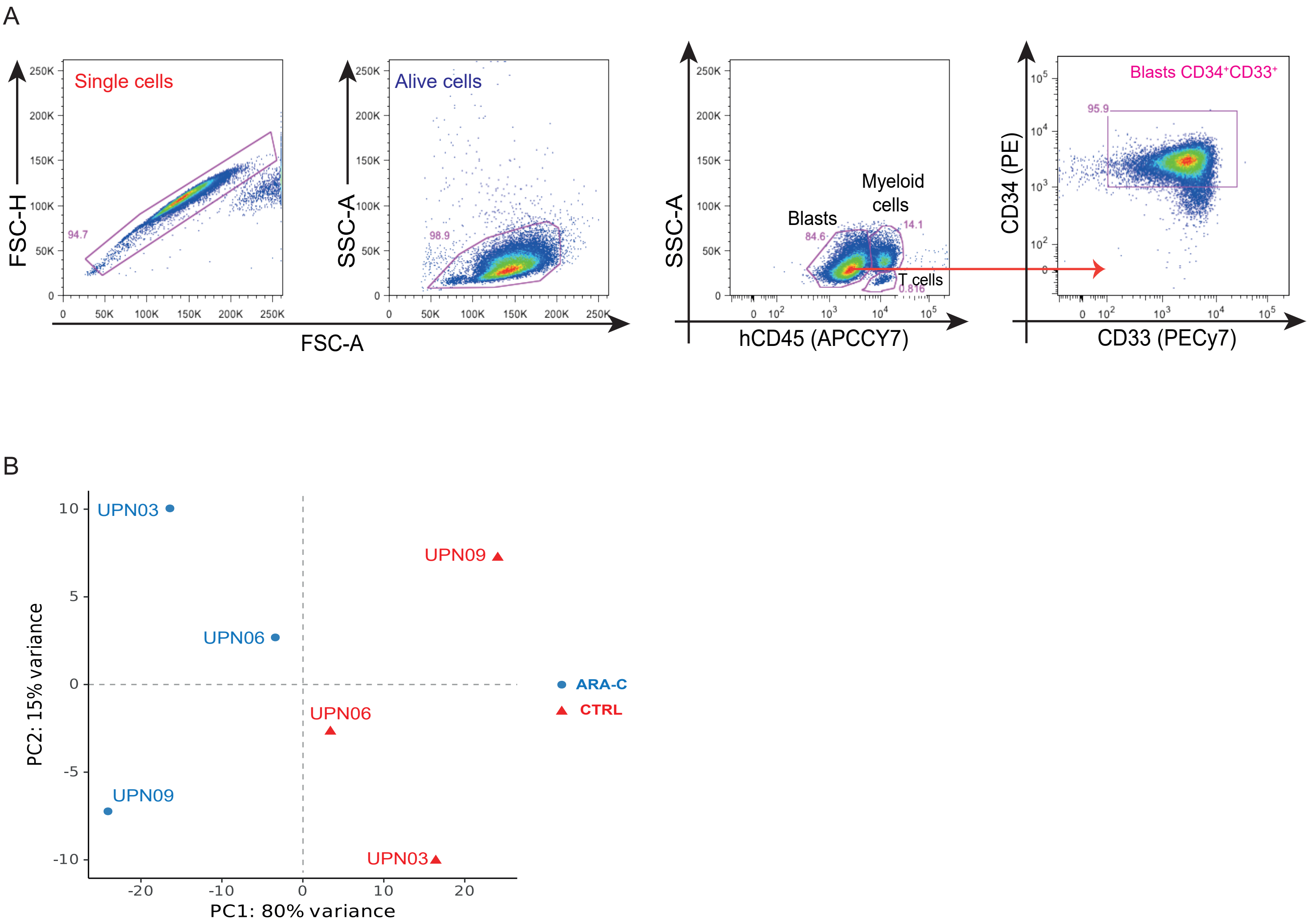

### Supplementary figure 2

Fig. S2 Senescence establishment in primary AML and cell lines

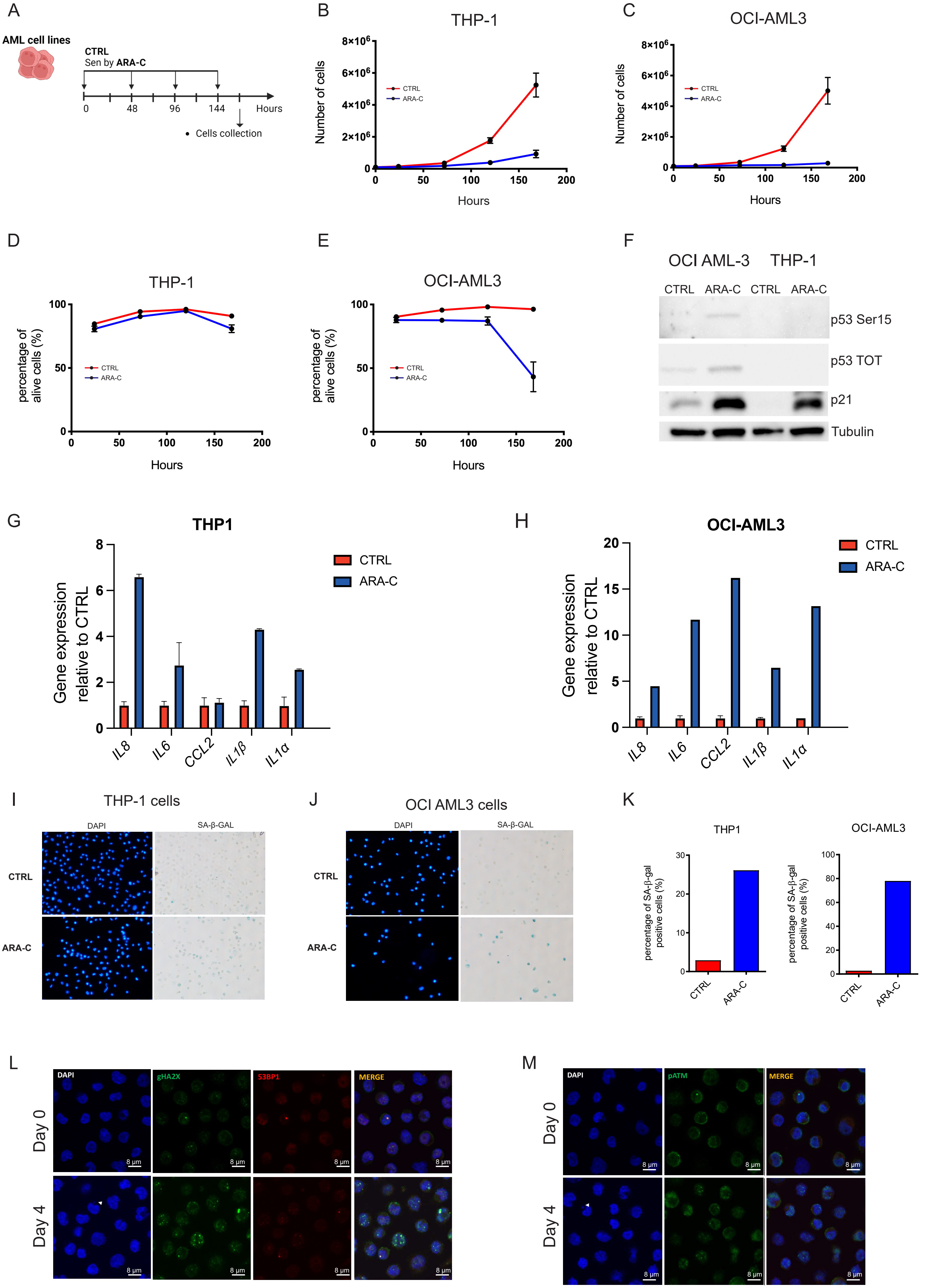

### Supplementary figure 4

Fig. S4 Chronic or acute effects of senescence-inducing treatments in AML cell lines

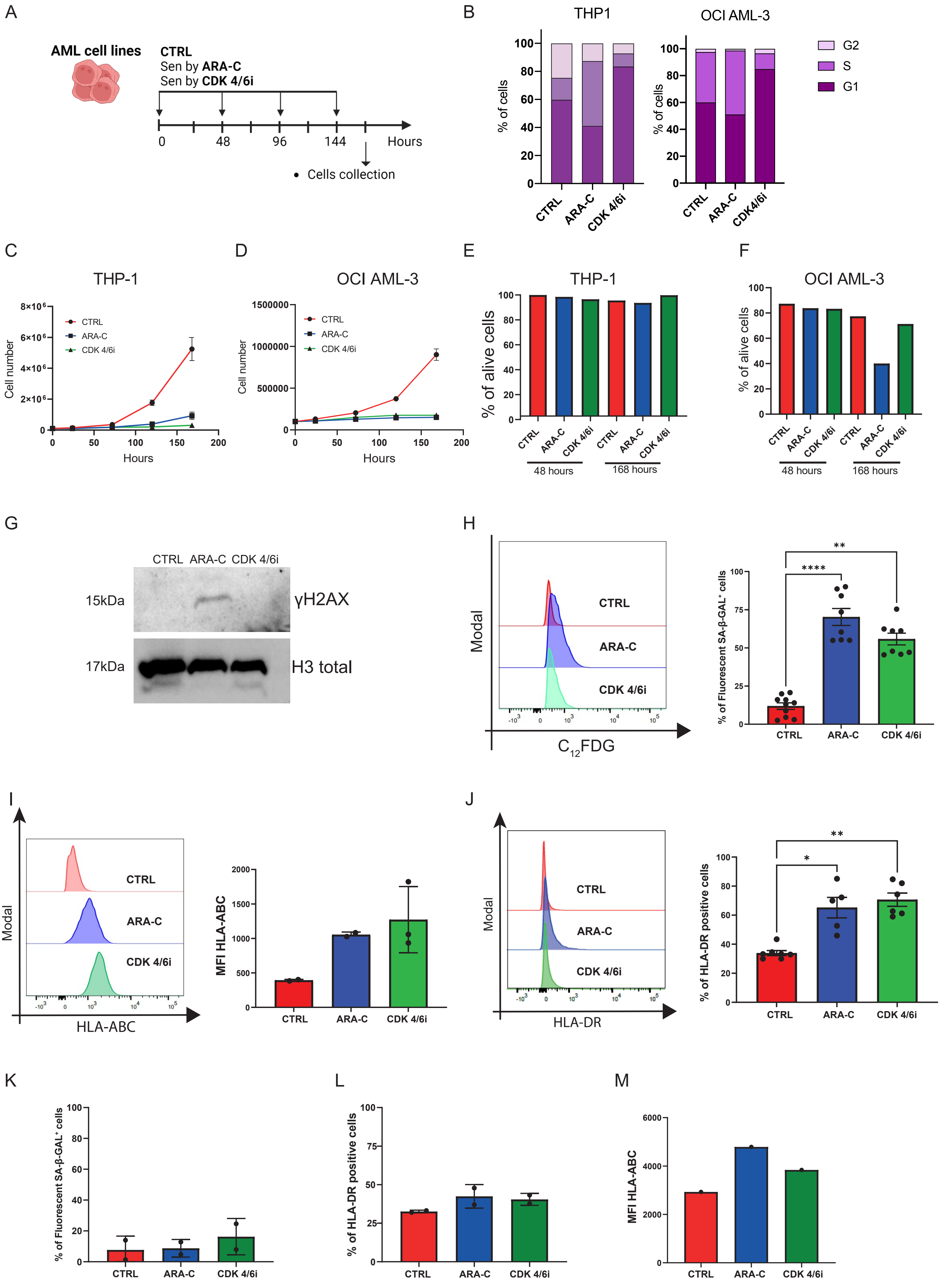

### Supplementary figure 5

Fig. S5 Analyses of senescence-induced T-cell activation

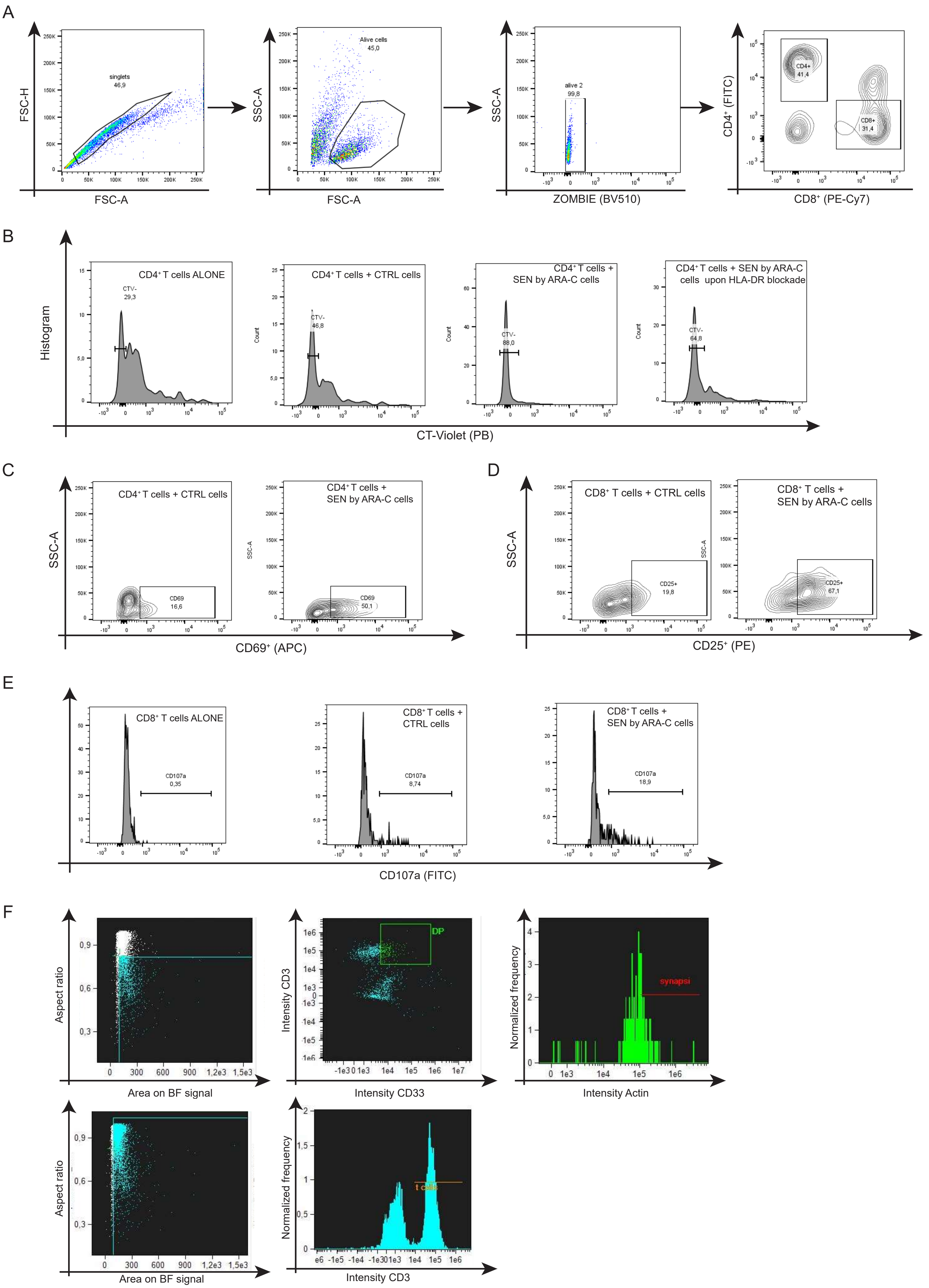
