## Supplementary figure 3 for "Therapy-induced senescence upregulates antigen presentation machinery and triggers anti-tumor immunity in Acute Myeloid Leukemia"

Fig. S3 Immune-related categories and HLA-class I or HLA-class II levels upon senescence induction in primary AML

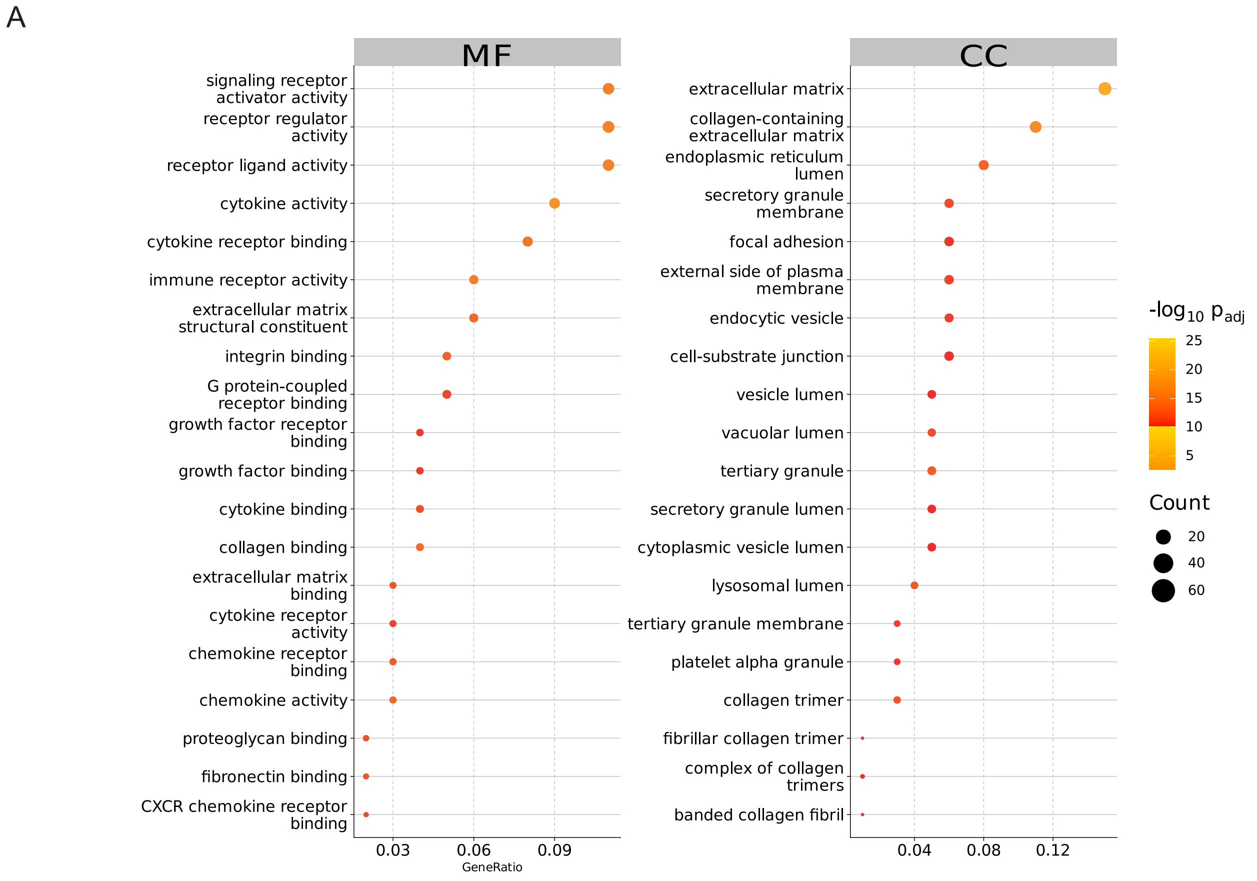


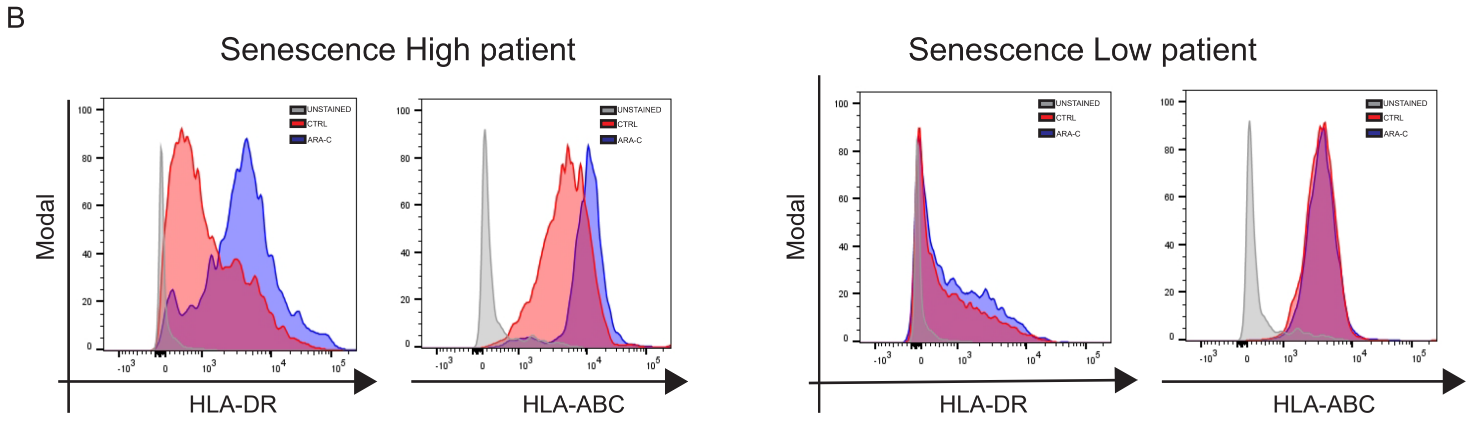
