## Supplementary figure 6 for "Therapy-induced senescence upregulates antigen presentation machinery and triggers anti-tumor immunity in Acute Myeloid Leukemia"

Fig. S6 Activation and exhaustion profiles of T cells from AML patients at diagnosis or 30 days post-chemotherapy

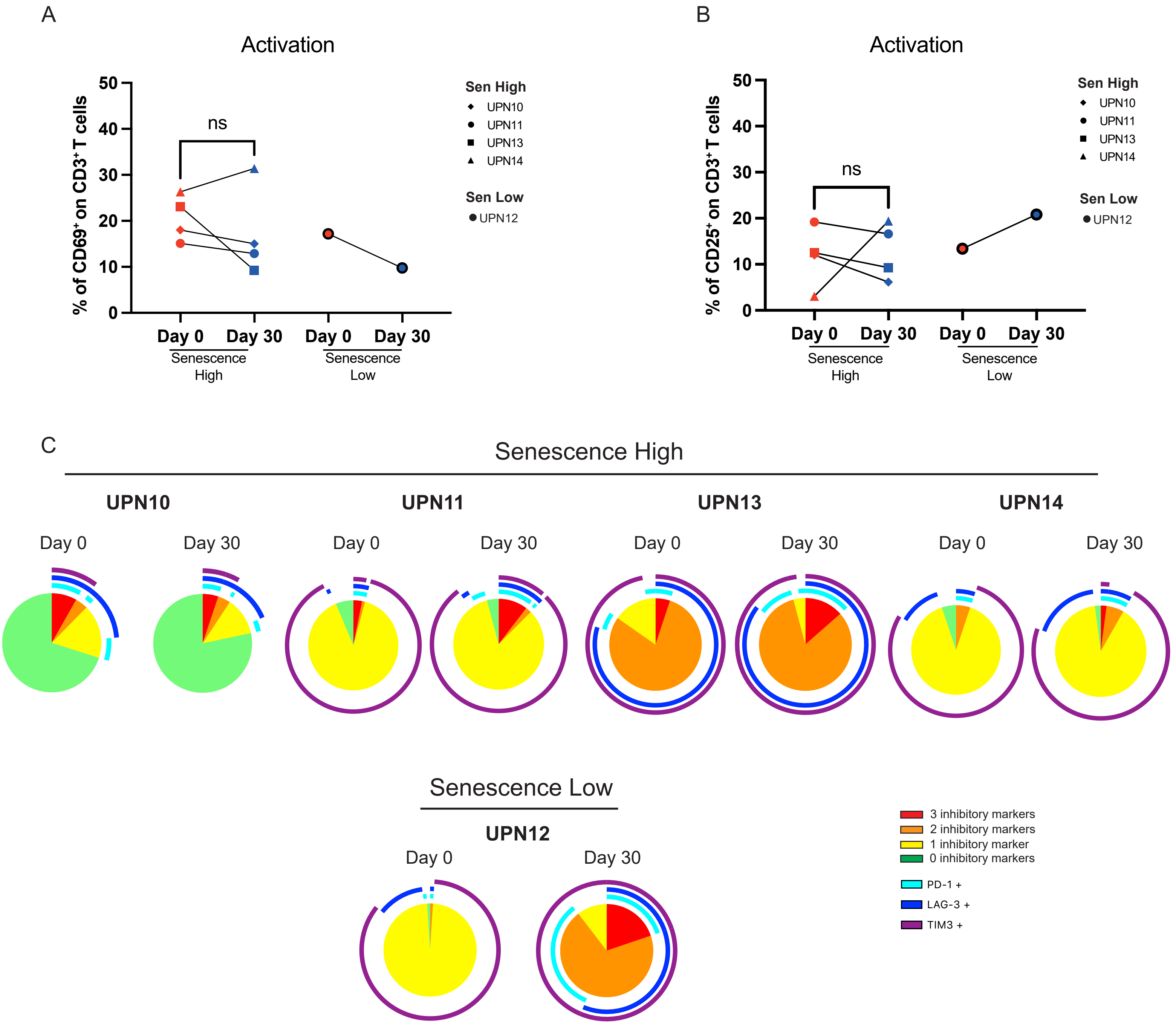
