## Supplementary table S1 for "Therapy-induced senescence upregulates antigen presentation machinery and triggers anti-tumor immunity in Acute Myeloid Leukemia"

| **Patient  Code** | **Sex** | **Age at  Diagnosis  (Y)** | **Cytogenetics** | **Molecular  Alterations*** | **Induction Chemotherapy** |
| --- | --- | --- | --- | --- | --- |
| **UPN 01** | M | 70 | normal | - | 3+7 |
| **UPN 02** | M | 41 | normal | FLT3-ITD | 3+7 |
| **UPN 03** | F | 60 | normal | FLT3-ITD NPM1mutA | 3+7 |
| **UPN 04** | M | 72 | t(10;13) (q24;q22) | FLT3-ITD NPM1mutA | 3+7 |
| **UPN 05** | M | 70 | normal | FLT3-ITD | 3+7 |
| **UPN 06** | M | 52 | Unknown | FLT3-ITD | 3+7 +FLT3 inhibitor |
| **UPN 07** | F | 70 | normal | - | 3+7 |
| **UPN 08** | M | 71 | del(15)(q?) | - | 3+7 |
| **UPN 09** | M | 48 | -Y,t(8;21)(q22;q22) | - | 3+7 |
| **UPN 10** | M | 43 | normal | NPM1mutA | ICE |
| **UPN 11** | F | 46 | inv(16)(p13.1q22) | - | 3+7 |
| **UPN 12** | M | 65 | normal | - | 3+7 |
| **UPN 13** | M | 44 | normal | NPM1mutA | 3+7 |
| **UPN 14** | F | 67 | der(5)t(5;?)(q31.3;?),  t(7;21)(q22;q22), inv(16)(p13.1q22) | - | 3+7 |
| **UPN 15** | M | 70 | +13 | - | 3+7 |
| **UPN 16** | F | 30 | t(11;19) (q23;p13) | TP53 | 3+7 |
| **UPN 17** | M | 62 | +11 | - | 3+7 |
| **UPN 18** | F | 35 | add(10)(p?15),-11,add(12)(p?13), -16,-16,-18,+3mar | - | 3+7 |
| **UPN 19** | M | 62 | normal | - | 3+7 |
| **UPN 20** | F | 66 | normal | FLT3-ITD NPM1mutA | 3+7 |
| **UPN 21** | F | 57 | 44,XX, der(2)t(1;2)(p?34;p?23), del(5)(q13q33), -7,  add(16)(q?1213), 17, 18, 21, +2mar | - | 3+7 |

*Mutational profiles included analyses for *NPM1*, *FLT3-ITD*, *FLT3-TDK*; *CEBP-alpha* and *TP53*. ICE: idarubicin, cytarabine and etoposide. *Ex-vivo* chemotherapy-treated patients: **Senescence High**: UPN06, UPN07, UPN08, UPN09, UPN10, UPN11, UPN13, UPN14, UPN15, UPN16. **Senescence Low**: UPN01, UPN02, UPN03, UPN04, UPN05, UPN12, UPN17. *In-vivo* chemotherapy-treated patients: UPN18, UPN19, UPN20, UPN21.
